# The neural cell adhesion molecule NrCAM regulates development of hypothalamic tanycytes

**DOI:** 10.1101/2021.12.15.472761

**Authors:** Alex Moore, Kavitha Chinnaiya, Dong Won Kim, Sarah Brown, Ian Stewart, Sarah Robins, Georgina Dowsett, Charlotte Muir, Marco Travaglio, Jo E. Lewis, Fran Ebling, Seth Blackshaw, Andrew Furley, Marysia Placzek

**Author notes:** Indicates equal contribution (Alex Moore, Kavitha Chinnaiya). Correspondence to, (M.P.).

## Abstract

Hypothalamic tanycytes are neural stem and progenitor cells, but little is known of how they are regulated. Here we provide evidence that the cell adhesion molecule, NrCAM, regulates tanycytes in the adult niche. NrCAM is strongly expressed in adult mouse tanycytes. Immunohistochemical and *in situ* hybridization analysis revealed that NrCAM loss of function leads to both a reduced number of tanycytes and reduced expression of tanycyte-specific cell markers, along with a small reduction in tyrosine hydroxylase-positive arcuate neurons. Similar analyses of NrCAM mutants at E16 identify few changes in gene expression or cell composition, indicating that NrCAM regulates tanycytes, rather than early embryonic hypothalamic development. Neurosphere and organotypic assays support the idea that NrCAM governs cellular homeostasis. Single-cell RNA sequencing (scRNA-Seq) shows that tanycyte-specific genes, including a number that are implicated in thyroid hormone metabolism, show reduced expression in the mutant mouse. However, the mild tanycyte depletion and loss of markers observed in NrCAM-deficient mice were associated with only a subtle metabolic phenotype.

## Introduction

The hypothalamus is the central regulator of organism-wide homeostasis and underpins numerous processes that support life, including feeding behaviour, energy balance, sexual behaviour and reproduction, aggression, and response to stress (Lechan and Toni, 2016). Situated around the third ventricle (3V) at the base of the forebrain, its neurons are found both within dense nuclei and more sparse areas that extend from the supraoptic region at the anterior border to the mammillary bodies at the posterior border. At the ventral surface of the mediobasal/tuberal hypothalamus, the median eminence (ME) and neurohypophysis/ posterior pituitary bulge ventrally out of the hypothalamic pial surface, forming critical interfaces between the brain and the peripheral circulation.

Tanycytes - specialised Nestin-positive radial glial-like cells whose soma line the lateral and ventral walls of the 3V - are key to the ability of the ME to support the two-way flow of information from body-to-brain and brain-to-body (Prevot et al., 2018; Rodríguez et al., 2019). Four distinct tanycyte subsets are recognised (α1-, α2-, β1- and β2-tanycytes) (Rodríguez et al., 2019) that have been classified according to dorso-ventral (D-V) position, morphology, and marker expression – as well as through single-cell RNA sequencing (scRNA-Seq). Further, in rodents, different tanycyte subsets have been classified according to function. Distinct tanycyte subsets have been shown to regulate hormone levels, act as nutrient and hormone sensors, and regulate body content – all of which support acute physiological changes (Samms et al., 2015; Lazutkaite et al., 2017; Müller-Fielitz et al., 2017; Farkas et al., 2020; Yoo et al., 2020; Rohrbach et al., 2021).

In addition, studies in mice show that tanycytes may regulate long-term changes to hypothalamic neural circuitry through their ability to act as stem and progenitor cells (Lee et al., 2012, 2014; Robins et al., 2013; Yoo et al., 2021). Lineage-tracing and genetic ablation studies *in vivo*, and neurospherogenic studies *ex vivo*, support the idea that α2-tanycytes are stem-like cells, and that other tanycyte subsets are progenitor-like cells (Robins et al., 2013; Yoo et al., 2021). While tanycytes can be experimentally forced into a neurogenic phase, the levels of tanycyte-derived neurogenesis are extremely low under baseline conditions in adult mouse, and the extent and physiological importance of tanycyte-derived neurogenesis in response to stimuli such as high-fat diet or psychological stress remains unclear (Yoo and Blackshaw, 2018). Further, as yet, little is known about the control of stem-like tanycytes. In other stem cell niches within the brain, cues provided by the local microenvironment ensure a fine balance that supports the sustained self-renewal of multipotent stem--like cells with the generation of progenitor cells and differentiated neurons (Silva-Vargas et al., 2013; Bjornsson et al., 2015). This balance allows continuous neurogenesis without depletion of the neural stem cell (NSC) pool and is tightly regulated. To date, it is unclear what factors support the maintenance of stem-like hypothalamic tanycyte cells.

Increasingly, studies have begun to focus on the role of cell adhesion molecules (CAMs) in regulating NSCs within their niche ((Hynes, 2002; Marthiens et al., 2010; Hu et al., 2017; Morante-Redolat and Porlan, 2019). CAMs from the cadherin, integrin, selectin, and immunoglobulin families are expressed within neurogenic regions of the adult brain (Chen et al., 2013), and members of the immunoglobulin superfamily of CAMs (Ig-SF CAMs) have been shown to play roles in the regulation of neural progenitors (Gennarini and Furley, 2017). However, while CAMs can regulate acute physiological functions in tanycytes (Messina and Giacobini, 2013; Parkash et al., 2015), no study has yet asked whether CAMs regulate the stem/progenitor-like behaviour of tanycytes.

The L1-like family of CAMs are a prominent subgroup of Ig-SF CAMs that play a wide range of roles in neuronal differentiation (Maness and Schachner, 2007). In addition, L1CAMs, including the family member NrCAM (NgCAM-related cell adhesion molecule/neuronal cell adhesion molecule), modulate neural cell proliferation. In mouse, loss of NrCAM leads to a reduction in cerebellar lobe size, while L1/NrCAM double mutants show major reductions in cerebellar size (Sakurai et al., 2001; Heyden et al., 2008; Zonta et al., 2011). A role for NrCAM in controlling cell proliferation is further implied from its involvement in tumorigenesis, where it has been linked to increased proliferation and motility in melanoma and colorectal cancers (Conacci-Sorrell et al., 2002, 2005; Chan et al., 2011).

Previous studies have suggested expression of NrCAM in the ventricular zone (VZ) of both the developing ((Edwards et al., 1990; Lustig et al., 2001; Noctor et al., 2001; Tamamaki et al., 2001; Sild and Ruthazer, 2011) and the adult mouse hypothalamus (Lein et al., 2007), although no detailed analysis has been described. Furthermore, both studies of NrCAM-null mice and genomic association studies have linked NrCAM with impaired sociability, addiction and autism, conditions associated with hypothalamic dysfunction (Ishiguro et al., 2006, 2014, 2019; Sakurai et al., 2006; Moy et al., 2009; Sakurai, 2012). As yet, though, no study has systematically analysed the expression or potential function of NrCAM in the hypothalamus.

Here we describe the cellular expression pattern of NrCAM in the hypothalamus of 8-10 week adult mice. NrCAM is detected in all four tanycyte subsets and in hypothalamic astrocytes. In hypothalamic neurospheres, NrCAM is coexpressed with hypothalamic stem/progenitor-like regional markers, indicating that it marks tanycyte stem/progenitor cells. Loss of NrCAM (NrCAM KO) leads to a small but significant loss of tanycytes and reduction in expression of tanycyte markers, a thinning of the VZ, and a modest but statistically significant reduction in TH-positive neurons in the postnatal period. Further, *ex vivo* assays show that tanycytes derived from NrCAM KO mice show decreased proliferation and generate smaller numbers of GFAP-positive astrocytes and TH-positive neurons. ScRNA-Seq of hypothalamic tissue from wildtype and NrCAM KO littermates confirms that NrCAM is highly expressed in tanycytes and astrocytes. In the tanycyte-cell fraction of NrCAM KO mice, tanycyte-specific genes, including a number that are implicated in thyroid hormone metabolism, show reduced expression, while astrocyte-specific genes show increased expression. In contrast, expression of astrocyte-specific markers are reduced in mutant astrocytes. We also detect altered expression of some GABAergic markers and *Trh* in hypothalamic neurons. NrCAM KO mice showed a small but significant reduction in body weight compared to wildtype littermates, which was associated with reduced food intake, but not with altered locomotor activity or energy expenditure. In summary, NrCAM is a novel surface marker of hypothalamic tanycytes and may regulate cellular homeostasis in and around the 3V. However, the mild tanycyte depletion and loss of markers observed in NrCAM KO mice were associated with only a subtle metabolic phenotype.

## Results

### Stem and progenitor cell marker distribution in the adult hypothalamus

We systematically analysed Nestin-positive tanycyte distribution over a 1000 μm length in and around the ME, corresponding approximately to Bregma position −0.9 to −2.2 in the intact skull. From anterior to posterior, tanycytes are first detected around the ventral-most VZ, anterior to the morphologically-distinct ME (a region we term the anterior ME (AME), corresponding to sections 67-68 in the Allen Brain Reference Atlas (ABRA)) (Fig 1A, A’). Over the length of the ME (corresponding to sections 69-73 in the ABRA) they increase in density and extend further dorsally (Fig 1B, B’). Tanycytes continue to line the 3V in the posterior ME (PME) (sections 74-75 in the ABRA) and a more posterior region adjacent to the periventricular nucleus (posterior part) (sections 76, 77 in the ABRA) (Fig 1C, D, C’, D’). At all positions, Nestin is detected strongly in β1/2 tanycytes, and more weakly in dorsal α1/2 tanycytes. Direct comparison of Nestin and GFAP – a marker of α2-tanycytes – confirms that GFAP-positive α2-tanycytes-previously implicated as stem-like cells - lie dorsal to β-tanycytes, and shows that GFAP-positive tanycytes are largely absent from the AME and ME (Fig 1E-H’).

**Figure 1.**
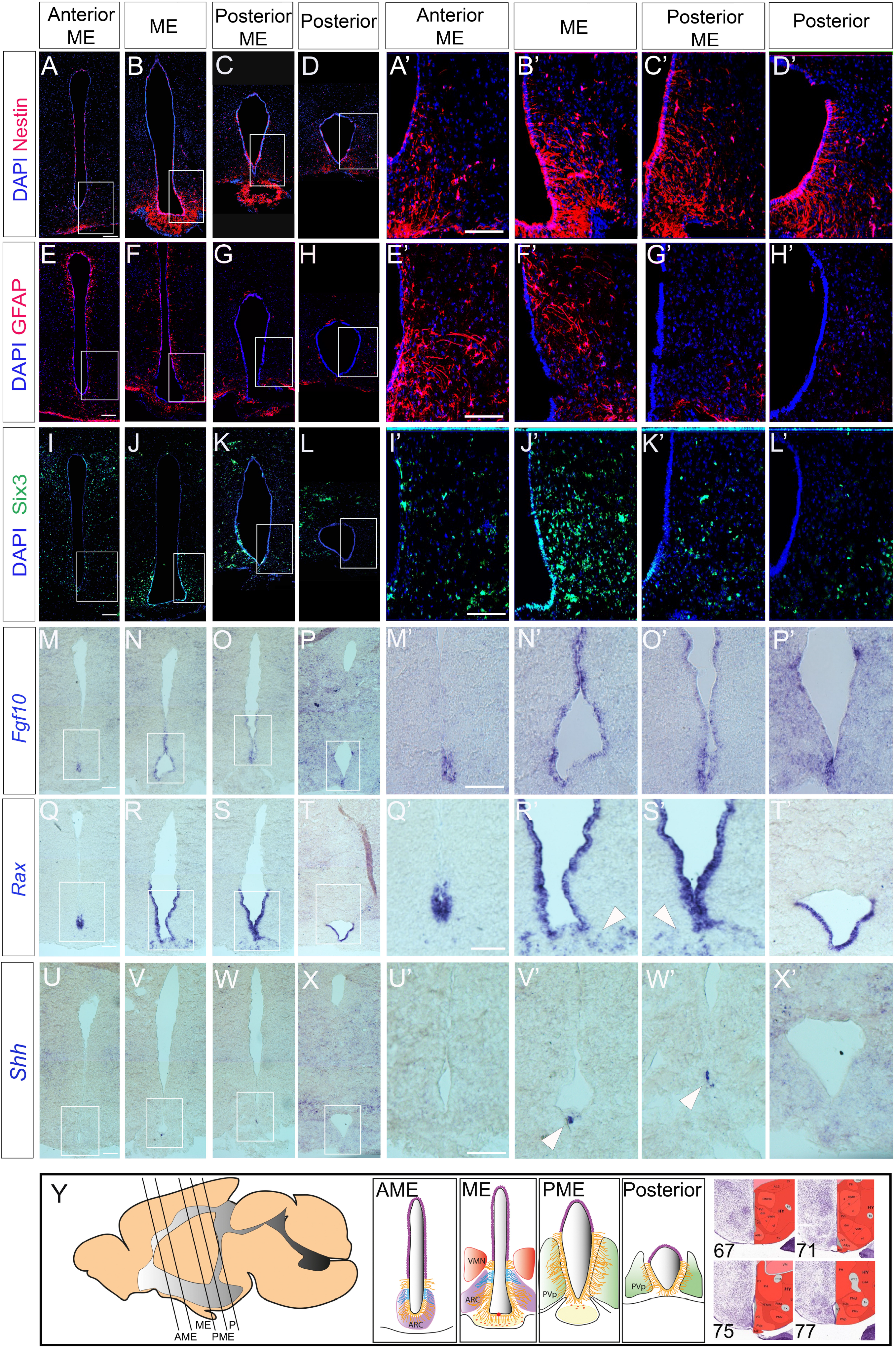
Stem and progenitor marker distribution in the adult hypothalamus. A-L’. Representative examples from consecutive serial adjacent coronal sections, from anterior to posterior, at the level of the AME, ME, PME and posterior tanycyte-rich hypothalamus, analysed for expression of Nestin, GFAP and Six3. Boxed regions in A-D, EH, I-L shown at high power in A’-D’, E’-H’, I’-L’. (n=5 mice; images from a single mouse) M-X’. Representative examples from consecutive serial adjacent coronal sections across the AME, ME, PME and posterior tanycyte-rich hypothalamus, analysed for expression of *Fgf10, Rax* and *Shh*. Boxed regions in M-P, Q-T, U-X shown at high power in M’-P’, Q’-T’, U’-X’. Arrowheads in R’,S’ point to *Rax*-expressing displaced tanycytes; arrowheads in V’, W” point to *Shh*-expressing VZ cells. (n= 3 mice; images from a single mouse) Y. Schematics showing approximate distribution of tanycytes. Sagittal schematic shows approximate section planes and positions of tanycytes along the A-P axis, in AME, ME, PME and posterior regions. Coronal schematics show extent of tanycytes (marked in yellow, apart from GFAP-positive tanycytes, marked in blue) along the D-V axis. Red dot shows position of *Shh*-positive VZ cells; orange dots show displaced *Rax*-positive tanycytes. Positions of the AME, ME, PME and posterior were judged relative to sections in the ABRA (right: ABRA section number shown), based on morphology and position of key nuclei. Scale bar: 100μm. Abbreviations: ARC, arcuate nucleus; AME, anterior to median eminence; ME, median eminence, PME, posterior median eminence; P, posterior tanycyte-rich hypothalamus; PvP, periventricular nucleus, posterior part; VMN, ventromedial nucleus.

We next asked how the gene expression profile of tanycytes compares to that of other known hypothalamic stem/progenitor markers. Direct comparison of Nestin and Six3 shows that in the VZ, Six3 is largely confined to ME/PME regions with little/no expression detected in AME or posterior regions (Fig. 1I-L’). *Fgf10* – implicated in the control of tanycyte proliferation (Robins et al., 2013) – is expressed in the AME, ME and PME, but is barely detected in the posterior hypothalamus. By contrast, *Rax* – a transcription factor necessary for tanycyte differentiation (Miranda-Angulo et al., 2014; Salvatierra et al., 2014) - is expressed throughout tanycyte-rich regions in the AME, ME, PME and posterior hypothalamus (Fig 1Q-T’). In addition, *Rax* is detected in ME cells beneath the VZ - previously described as displaced tanycytes (Miranda-Angulo et al., 2014) (Fig 1R’,S’ arrowheads). In the embryo, Six3, Rax and Fgf10 all modulate the activity of the signalling ligand Shh to direct differentiation of anterior tuberal progenitor cells (Kano et al., 2019; Farkas et al., 2020; Placzek et al., 2020). This, and the overlap of *Six3/Rax/Fgf10* in the adult, prompted us to examine expression of *Shh* in the adult hypothalamus. *Shh* was detected, but confined to a tiny subset of cells, occupying the midline in the ME/PME (Fig 1U-W’). Together these analyses show a previously undescribed profile of tanycyte distribution along the A-P axis (schematised in Fig 1Y).

### NrCAM is expressed on tanycytes

We next examined the profile of NrCAM. Double-labelling of consecutive sections with Nestin and NrCAM reveals that NrCAM is expressed in tanycytes throughout the AME, ME, PME and posterior hypothalamus (Fig.2A-D). At all positions, NrCAM is detected strongly on all tanycyte cell bodies throughout the VZ (Fig 2A-E, G-J) and more weakly on tanycyte processes (Fig 2A-E, H-J), which are fasciculated (Fig 2F). While there is some variation in expression levels of Nestin and NrCAM in different tanycyte subsets (Fig 2H, I), quantitative analyses of high power views suggest similar numbers of Nestin-positive and NrCAM-positive tanycytes (Fig 2K).

**Figure 2.**
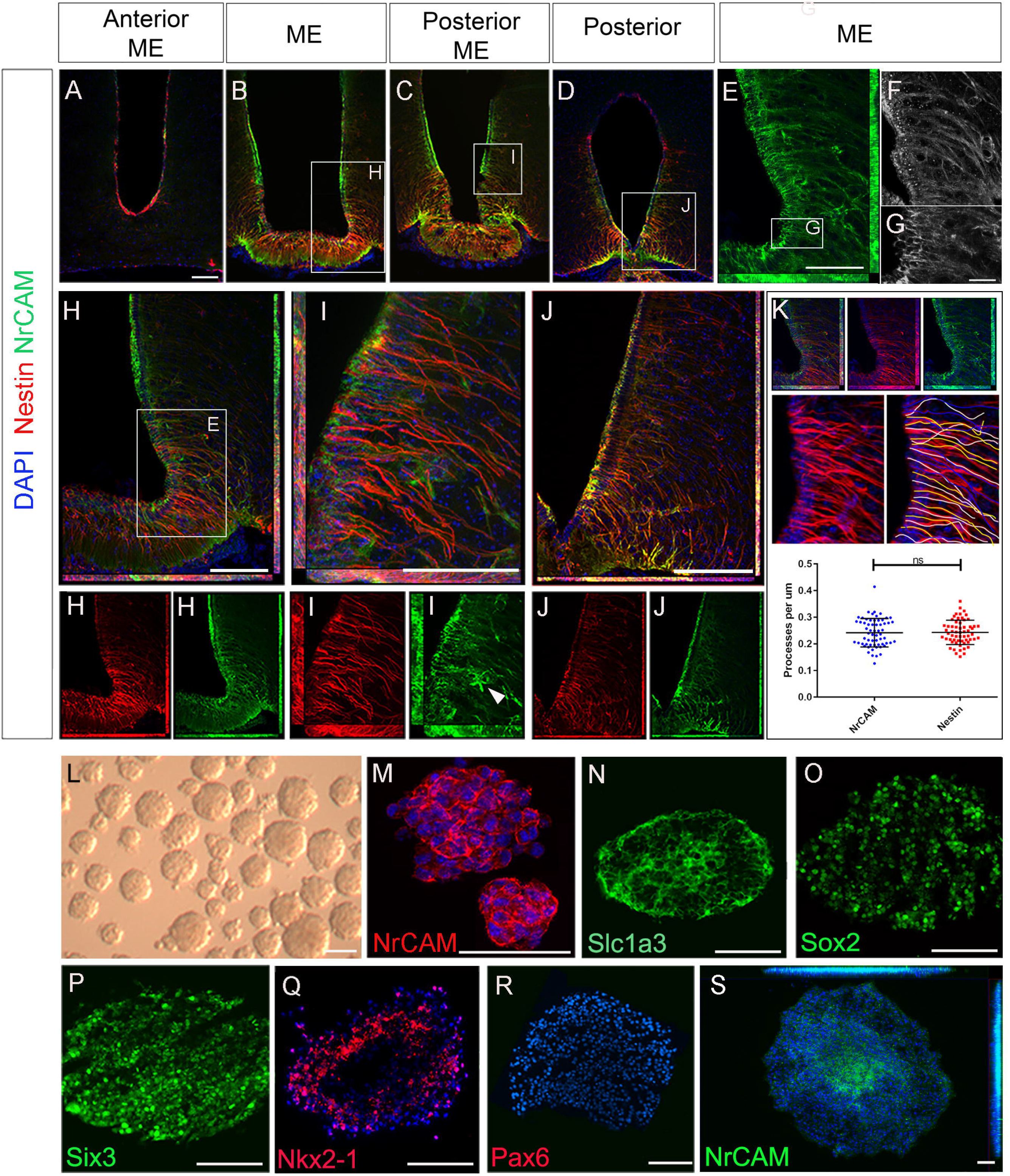
NrCAM is expressed on self-renewing hypothalamic tanycytes. A-D, H-J. Representative examples from consecutive serial coronal sections at the level of the AME, ME, PME and posterior tanycyte-rich hypothalamus, analysed for expression of Nestin and NrCAM. Boxed regions in B-D shown at high power in (H-J) as double or single channel views. Arrowhead in I (single channel green) points to an NrCAM-positive astrocyte. E. High power view of boxed region in (H). Single channel view shows NrCAM labelling F. High power view shows NrCAM labels fasciculated tanycytes G. High power view of the boxed region in (E) shows high expression of NrCAM on tanycyte cell bodies. K. Quantification of Nestin and NrCAM-positive process density. Sections through the hypothalamus (example shown in top row) were imaged at 40x and individual tanycytes traced (middle row). There is no significant difference across the entire hypothalamus (P<0.0001; unpaired t-test) (bottom row; n=3 mice). L-S. 7^th^ passage neurospheres derived from VZ around 3V, analysed in brightfield (L) or after immunohistochemical analyses (M-S) after culture under non-differentiation (M-R) or differentiation (S) conditions (n=15-20 neurospheres/condition). Scale bars: (A-E, H-J, L): 100μm; (G): 45μm; (M-S): 50μm

The particularly strong expression of NrCAM on α2 tanycyte cell bodies prompted us to explore whether NrCAM marks these cells as they undergo self-renewal. To do so we took advantage of previous work, which had shown that stem-like α2 tanycytes can be cultured as neurospheres, up to and beyond the seventh passage (Robins et al., 2013). Using this assay, we analysed 7^th^ passage neurospheres, cultured under non-differentiated conditions (Fig 2L). Neurospheres expressed NrCAM, which was detected on every cell (Fig 2M), as were the neural stem-like markers Slc1a3 and Sox2 (Fig 2N, O). At the same time, neurospheres expressed the hypothalamic stem/progenitor-like regional markers, Six3 and Nkx2-1 but did not express Pax6 or Nkx6-1 - markers of non-hypothalamic progenitor subtypes (Fig 2P-R). Expression of Nkx2-1 and Six3 was specific to hypothalamic-derived neurospheres and was not detected on neurospheres obtained from the subventricular zone (SVZ) or spinal cord ependymal cells (Supp Fig. S1). Under differentiation conditions, NrCAM became restricted to central-most portions of neurospheres (Fig 2S). Together this suggests that NrCAM marks tanycyte stem/progenitor cells.

### Loss of NrCAM depletes hypothalamic tanycytes postnatally

To determine if loss of NrCAM leads to a reduction in tanycyte number *in vivo*, we compared tanycyte process distribution and tanycyte progenitor marker expression in week 8-10 NrCAM KO mice - in which NrCAM protein cannot be detected (Supp Fig S2A, B) (Sakurai et al., 2001) - and wild-type littermates. Analysis of Nestin expression across the hypothalamus shows that fewer processes are seen in the AME, ME, PME, and P regions in NrCAM KO animals in comparison to WT animals (Fig 3A, B). Quantification of process density (analysis restricted to regions harbouring β1- and α2-tanycytes) showed a statistically significant decrease in process density across the whole hypothalamus (Fig 3K); consistently, each subregion showed an ~20% decrease in the number of Nestinpositive processes (Fig 3L). Analysis of GnRH, which decorates tanycyte processes, likewise points to a decrease in density of tanycyte processes (Supp Fig S2C, D). Analysis of the α2-tanycyte region, at the level of the ME (boxed region in Fig 3A, B) showed that the reduction in Nestin is accompanied by a significant thinning of the VZ (Fig 3A’, B’, M) and a marked reduction in GFAP-positive process density (Fig 3A’, B’, N). Analysis of Six3 expression confirms the thinner VZ (Fig 3C, D, O). At the same time, *Rax* and *Fgf10* are expressed more weakly in NrCAM KO mice, and show modest reductions in the size of their expression domain, compared to wildtype littermates (Fig 3E-H, P-S), while *Shh*-expressing cells are barely discernible (Supp Fig 2F, H’). This suggests that loss of NrCAM leads to a reduction in the generation and/or maintenance of tanycytes. Changes are not limited to ventricular cells, and we also note a significant reduction in the number of displaced *Rax*-positive tanycytes cells in the ME parenchyma (Fig 3E’, F’ arrowheads; Fig. 3T) (Miranda-Angulo et al., 2014).

**Figure 3.**
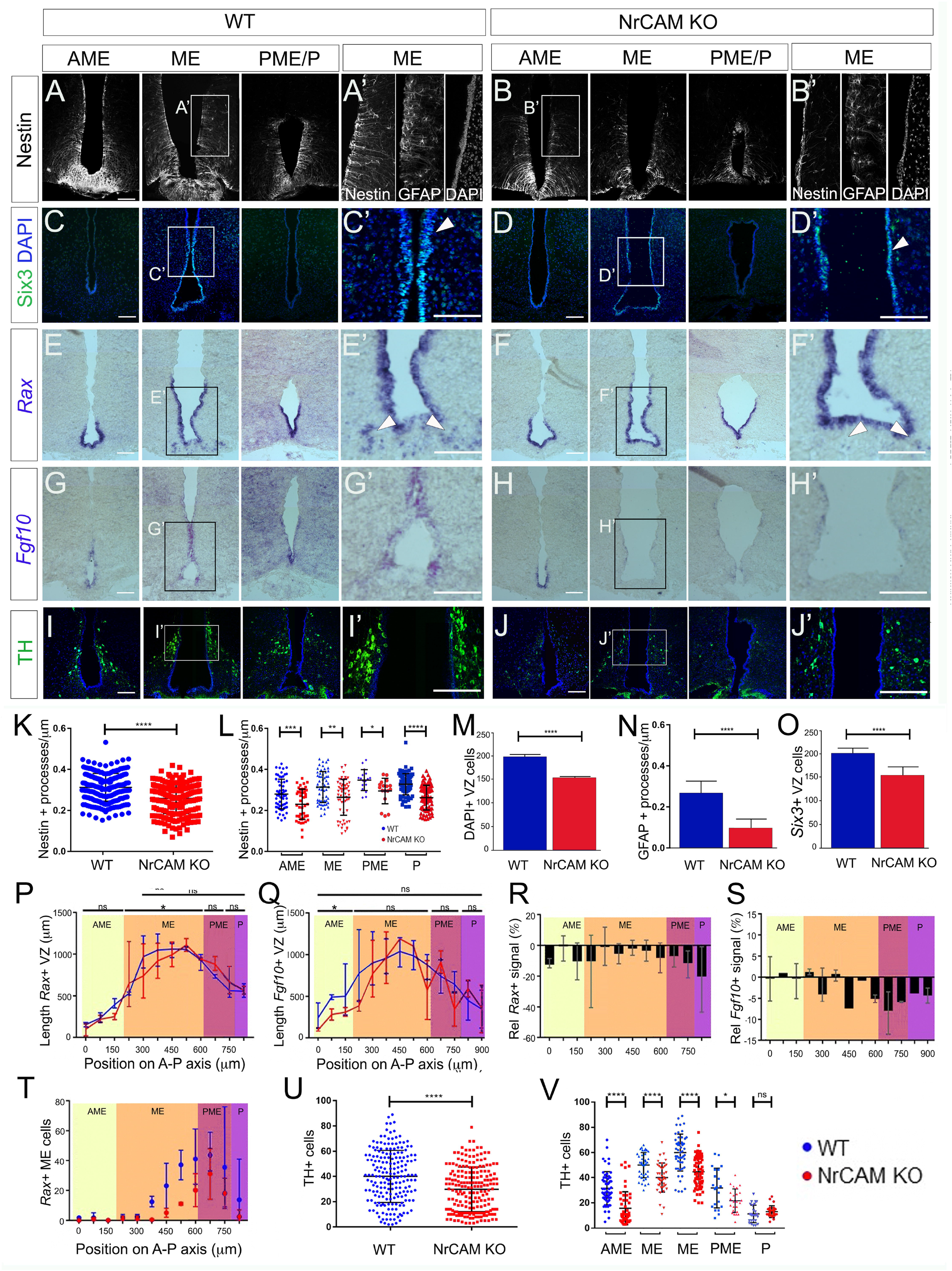
Reduced tanycytes and TH neurons in the NrCAM KO adult. A-J. Representative coronal sections through AME, ME and PME/P regions of the hypothalamus of wildtype (A, C, E, G, I) or NrCAM KO mice (B, D, F, H, J). (A’-J’) show high power views of boxed regions in (A-J). Sections were immunolabelled to detect Nestin (A, B), Six3 (C, D), TH (I, J), or Nestin/GFAP (A’,B’: high power views of boxed regions in (A, B), or analysed by *in situ* hybridisation to detect *Rax* and *Fgf10*. Arrowheads in (C’, D’) point to Six3-positive VZ cells and in (E’, F’) point to *Rax*-positive displaced tanycyte cells (n=8 mice/genotype: analysed for Nestin and TH (n=3); *Rax*, *Fgf10* (n=3); Six3 (n=2); images (A, I) (B, J), (E, G), (F, H) show serial consecutive sections through single mice. Scale bars: 100μm. K-V. Quantitative analyses in wildtype and NrCAM KO mice. (K, L) There is a significant reduction in Nestin-positive β1- and α2-tanycyte density in NrCAM KO mice across the entire hypothalamus (K) (P<0.0001; unpaired t-test) and across each subregion (L) (AME p=0.0007; ME p=0.0013; PME p=0.0105; p<0.0001; unpaired t-test). Each icon represents a single measurement (n=3 mice/genotype). Bars show SD. (M-O) There is a significant reduction in the number of cells lining the VZ (M), in GFAP-positive tanycyte density (N), and in Six3-positive nuclei in NrCAM KO mice (O) (p<0.0001; unpaired t-test) each. Bars show SE. (P-T) Quantitative analyses of *Rax* and *Fgf10*. Analysis was performed at 75μ intervals across AME, ME, PME and P subregions. Each plotted value is the mean of three biological replicates; bars show range. (P, Q): Lengths of *Rax*- and *Fgf10*-expressing domains in wildtype (blue) and NrCAM KO (red) mice and (R, S) percentage change in *Rax* and *Fgf10* signal strength in NrCAM KO relative to wildtype. *Rax*-expressing VZ was significantly longer in the ME subregion of wildtype mice (P) (Wilcoxon signed rank test; p=0.0273). *Fgf10*-expressing VZ was significantly longer in the AME subregion of wildtype mice (Q) (Wilcoxon signed rank test; p=0.0273). Relative intensity of *Rax* and *Fgf10* was reduced in NrCAM KO compared to wildtype mice at most levels (R, S). (T) Significantly fewer *Rax*-positive displaced tancytes were observed in the NrCAM KO compared to wildtype mice (Wilcoxon signed rank test; p=0.0010). (U, V) There is a significant reduction in TH-positive cells in NrCAM KO mice across the entire hypothalamus (U) (p<0.0001; unpaired t-test with Welch’s correction) and across subregions (V) (AME p<0.0001; ME p<0.0001; PME p=0.0158; unpaired t-test with Welch’s correction). No significant difference was seen in the P subregion where ARC TH^+^ cells are least populous (p=0.2865).). Each icon represents a single measurement. Bars show SD. (n=3 mice/genotype; 30 sections analysed/mouse for Nestin and TH; 10-12 sections analysed/mouse for Six3, *Rax*, *Fgf10*).

Previous studies have suggested that tanycytes can give rise to NPY-expressing neurons in the arcuate nucleus (Arc) (Haan et al., 2013; Yoo et al., 2021) and we therefore asked if the reduction in tanycytes in NrCAM KO mice is accompanied by a reduction in NPY neurons. Immunohistochemical analysis shows no change in area or intensity of NPY neurons in NrCAM KO vs wildtype littermates (Supp Fig S2E, G)). However, we detect a small but significant decrease in TH-positive A12 Arc neurons in NrCAM KO vs wildtype littermates (Fig 2I-J, U-V). The decrease in TH-positive neurons appears to be specific to the A12 population in the Arc region of the hypothalamus, as no significant difference was observed in the number of A13 TH-positive neurons of the zona incerta (Romanov et al., 2017) (Supp Fig S3).

Although these changes are consistent with the idea that NrCAM might play a role in cellular homeostasis within the hypothalamus, they could reflect a developmental phenotype: previous studies have shown that NrCAM is expressed in the hypothalamus in the E14 embryonic mouse (Lustig et al., 2001; Kim et al., 2020). Analysis of E16 mice, however, showed only a small decrease in density of radial glial processes, around the level of the emerging ME (Fig 4A, B, K; Supp Fig S4) and no difference in expression of *Rax*, Six3 (Fig 4C-F, L, M, P), *Fgf10*, or *Shh* (Fig 4N, O; Supp Fig S4). Lhx2, which is expressed at high levels in the cell bodies of α-tanycytes, shows a small decrease in the region of the ME, but in other regions, is expressed similarly in the two genotypes (Fig 4G, H, Q), while TH-positive neurons in the region of the developing Arc were increased in NrCAM KO mice, relative to wildtype littermates (Fig 4R, Fig S4; analysed at E18). The changes detected in 8-10 week mice therefore likely reflect an effect of NrCAM in the postnatal period.

**Figure 4.**
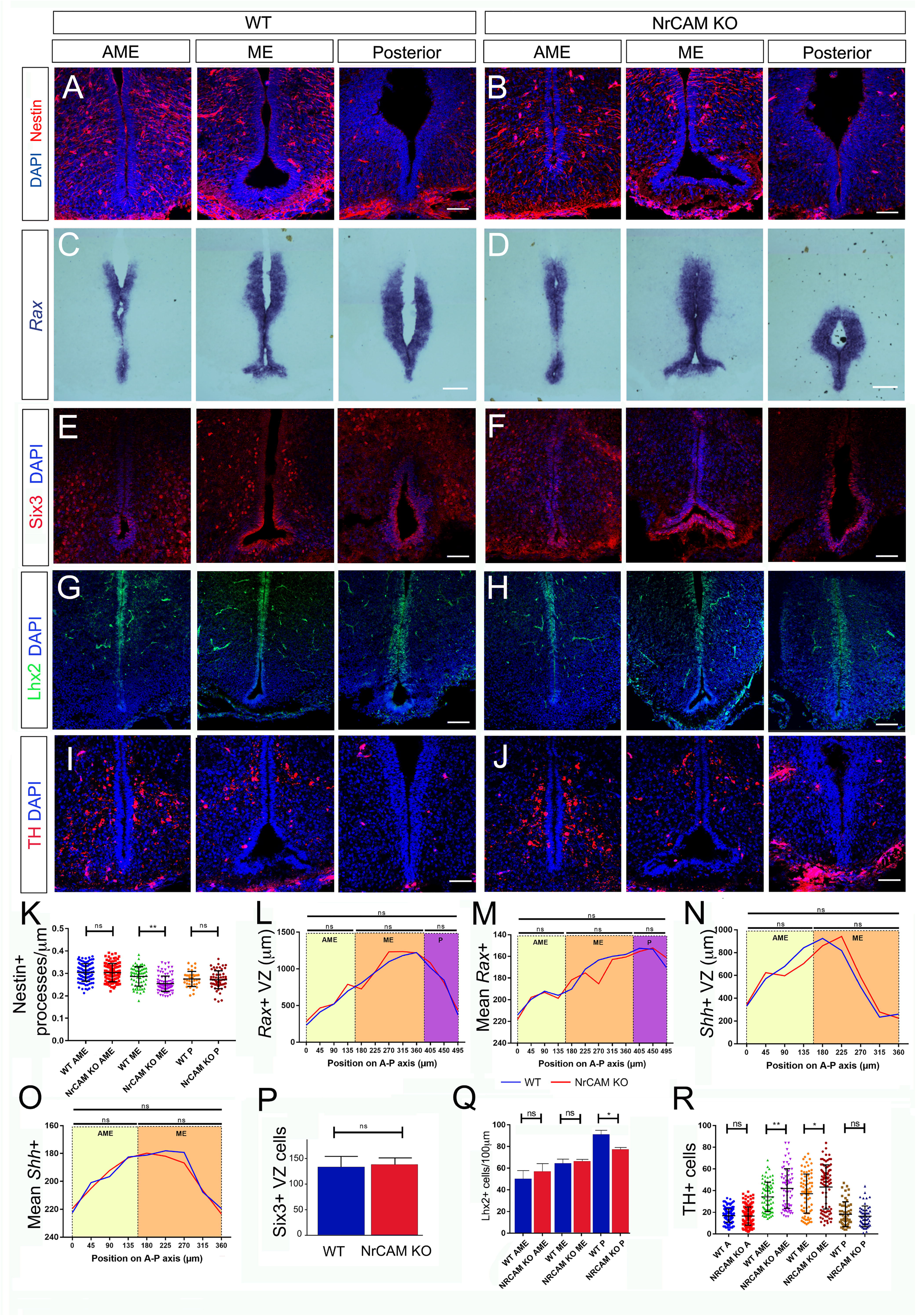
Comparison of tanycytes and TH neurons in the embryo. A-H. Representative serial coronal sections through AME, ME and posterior regions of the hypothalamus of E16 wildtype (A, C, E, G) or NrCAM KO mice (B, D, F, H) analysed by immunohistochemistry to detect Nestin (A, B), SIx3 (E, F), Lhx2 (G, H), or by chromogenic *in situ* hybridisation to detect *Rax* (C, D). (n=8 mice/genotype, analysed for Nestin (n=3), Six3, Lhx2 (n=2) or *Rax* (n=3). I, J. Representative serial coronal sections through AME, ME and posterior regions of the hypothalamus of E18 wildtype (I) or NrCAM KO mice (J) analysed by immunohistochemistry to detect TH. ((n=3 mice/genotype) Scale bars: 100μm. K-R. Quantitative analyses in wildtype and NrCAM KO mice. (K) There is only a small reduction in Nestin-positive β1- and α2-tanycyte density in NrCAM KO mice across the ME region (p<0.0200; unpaired t-test). Each icon represents a single measurement (n=3 mice/genotype). Bars show SD. (L-Q) There is no significant reduction in the lengths or relative intensities of *Rax*- and *Shh*-expressing domain (L-O), nor in the number of Six3+ VZ cells (P) in wildtype (blue) and NrCAM KO (red) mice. (Q) There is only a small reduction in the number of Lhx2+ VZ cells in NrCAM KO mice across the ME region (p=0.0052; unpaired t-test). (R) There is a small increase in TH-positive cells in NrCAM KO mice in the ME/AME (AME p=0.0044; ME p=0.0390) (10-12 sections analysed/mouse/genotype/marker)

### Decreased proliferation/differentiation of NrCAM-derived VZ/SVZ tissues

To directly assay the function of tanycytes in NrCAM KO mice, we analysed neurospherogenic potential of VZ cells harbouring β2 and α2 tanycytes. NrCAM KO hypothalamic tissue generated primary neurospheres that could be passaged. The number of secondary neurospheres formed from NrCAM KO was significantly higher than wildtype but then declined. By passage 7, the number of NrCAM KO neurospheres was significantly lower than wildtype neurospheres (Fig 5A). At the same time, cell concentration was significantly reduced in NrCAM KO neurospheres compared to wildtype (Fig 5B). These observations support the hypothesis that there are fewer stem/progenitor cells in NrCAM KO mice.

**Fig. 5.**
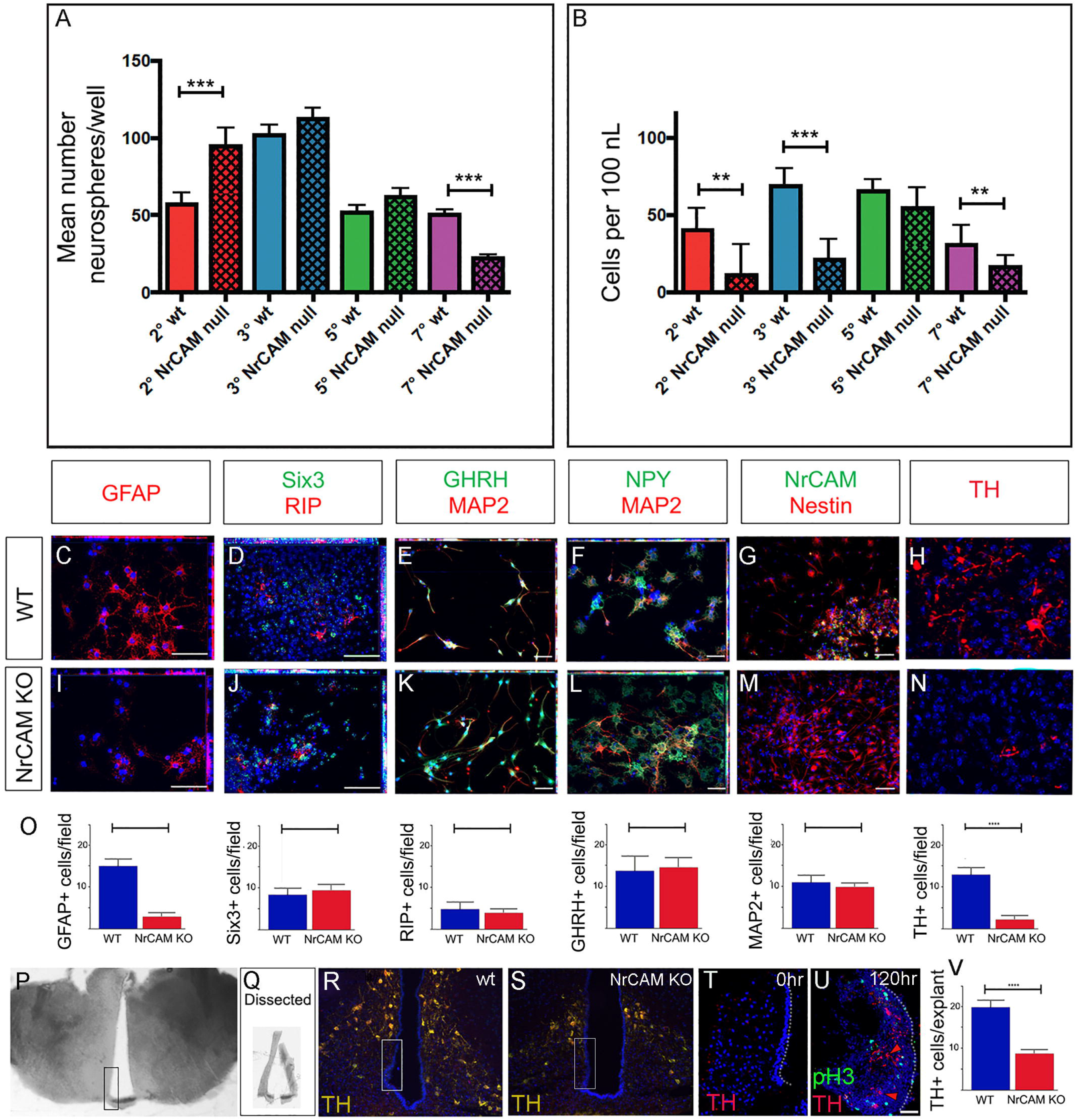
Decreased proliferation/differentiation of NrCAM-derived VZ/SVZ tissues. A. Average number neurospheres/well number in wildtype and NrCAM KO mice. The number of neurospheres from NrCAM null mice is significantly increased at passage 2 (p=0.0006) and is significantly decreased at passage 7 (p=<0.0001). Error bars show SE (n=6-12 wells; 3 mice each). B. The number of cells per ml cultured from wild type and NrCAM KO mice (calculated by dissociation of neurospheres from 12 wells). Neurospheres from NrCAM KO have a reduced number of cells per ml than wild type at passage 2, 3 and 7. Error bars show SE C-O. Fields of view (C-N) of neurospheres from passage 6, subject to differentiation conditions and analysed by immunohistochemistry after 7 days (C-G, I-M) or 14 days (H, N). Neurospheres from wildtype (C-H) and NrCAM KO littermates (I-N) show a similar range of differentiated cells. Quantitative analyses (O) show significantly fewer GFAP-positive astrocytes (unpaired t-test, p<0.0001) and significantly fewer TH-positive neurons (unpaired t-test, p<0.0001). (n=3 replicates; 20 fields of view total). P-V. Dissection and culture of 10 week hypothalamic tanycytes. (P) A 100μm thick slice through the region of the ME. Boxed region shows dissected area. (Q) Examples of dissected hypothalamic VZ. (R,S) Boxed regions show approximate position of dissected VZ in wildtype and NrCAM KO mice. (T) Representative example of a 15μm section through a VZ explant at t=0hr, immunolabelled with anti-TH. No TH-positive cells are detected. (U) Representative example of a 15μm section through a VZ explant at t=120hr, immunolabelled with anti-TH and anti-phosH3. (U) Quantitative analyses show significantly fewer TH-positive neurons after culture of VZ explants from NRCAM KO mice (unpaired t-test, p<0.0001). (n=10 explants/genotype, from 3 mice). Bars show SE. Scale bars: 50μm

To determine if neurospheres from NrCAM KO mice showed limited differentiation capacity, we subjected passage 6 neurospheres derived from NrCAM KO and wildtype littermates to differentiation conditions. As previously demonstrated (Robins et al 2013), wildtype hypothalamic neurospheres can give rise to both glial-like derivatives (GFAP-positive and RIP-positive cells) (Fig 5C, D) and hypothalamic neurons (GHRH, NPY, TH) (Fig 5E, F, H). Similar cells were detected in neurospheres derived from NrCAM KO mice (Fig 5I-N), and, as in wildtype tissue, the neurospheres retained some progenitor-like hypothalamic markers (Six3). Generally, similar numbers of cells were detected in the two genotypes; however, fewer GFAP-positive astrocytes and fewer TH-positive neurons were detected in NrCAM KO neurospheres compared to wildtype (Fig 5C, H, I, N, O). These analyses show that astrocytes and TH neurons can be generated from postnatal tanycytes as previously reported (Robins et al., 2013; Yoo et al., 2021), and suggest that loss of NrCAM reduces this efficacy.

Neurospheres, however, are cultured over a lengthy period, and so may be subject to culture artefacts. We therefore established an acute assay, to ask if we could see evidence for the de novo generation of TH-positive Arc neurons. VZ/SVZ tissue from the region of the ME was micro-dissected (Fig 5P, Q). We aimed to dissect tissue containing α2 and β-tanycytes, and avoid more dorsal regions that would likely be contaminated with TH-positive neurons (Fig 5R, S). Accuracy of dissection/avoidance of TH-positive neurons was confirmed through immediate analysis of a subset of explants (Fig 5T). In parallel, a subset of explants was cultured. In cultured explants from both wildtype and Nr CAM null mice, we detected phosH3-positive proliferating cells and TH-positive neurons (Fig 5U). However, we detected significantly fewer TH-positive neurons in NrCAM KO tissue, compared to wildtype (Fig 5V).

### scRNA comparison of NrCAM and wildtype hypothalamic tissue

To further identify molecular changes between wildtype and NrCAM KO, we performed scRNA-Seq on adult hypothalamus on both genotypes. This approach confirmed that *NrCAM* expression is detected in tanycytes and astrocytes, but also detected expression in neurons and oligodendrocyte-precursor cells (OPC) (Supp Fig S5). We then sub-setted tanycytes for further analysis, clustering these into α1-, α2-, β1- and β2 subsets based on previously known molecular markers (Fig 6A, B) (Yoo et al., 2021).

**Fig. 6.**
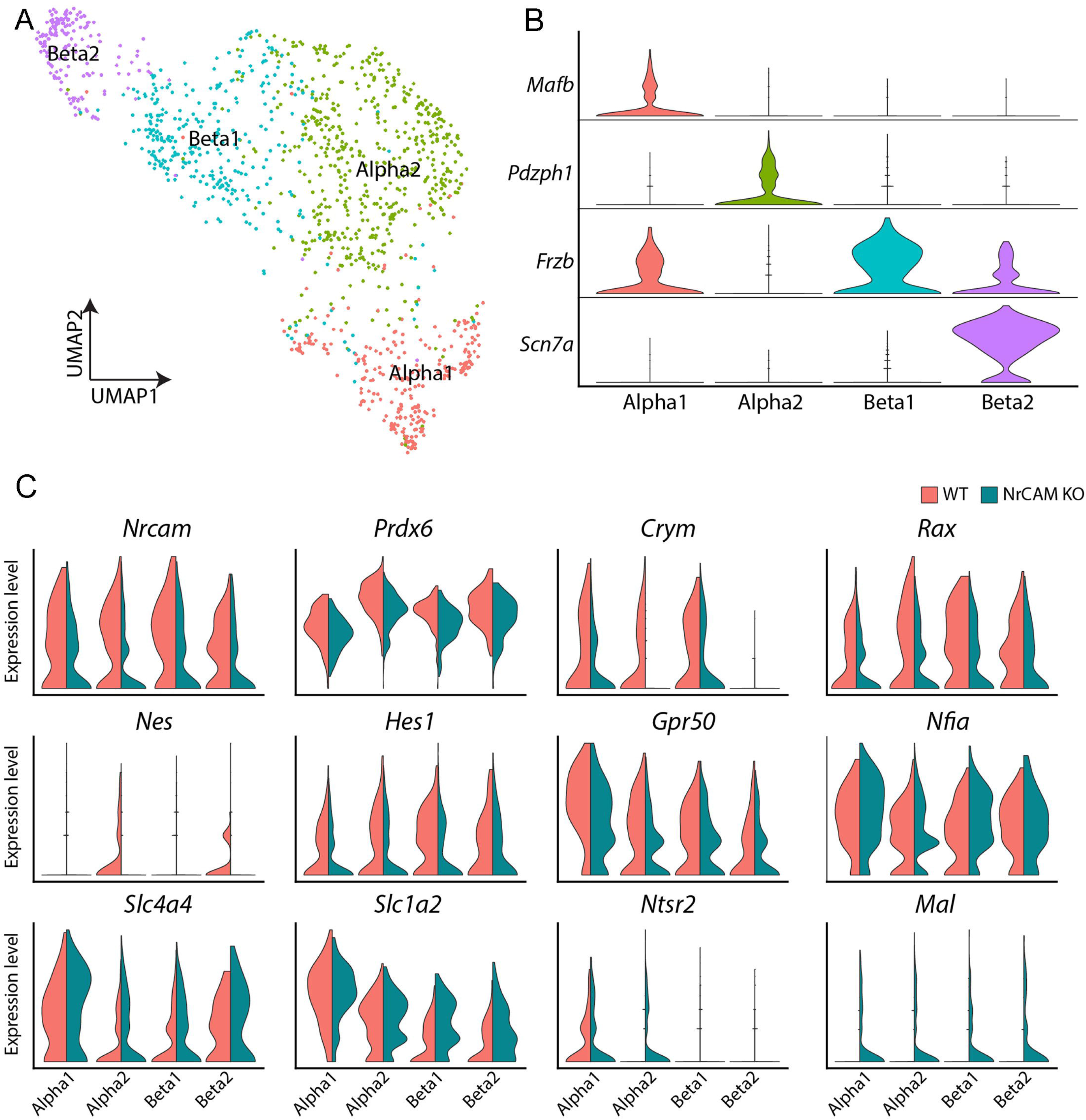
scRNA seq of hypothalamus from wildtype and NrCAM KO mice. A, B. UMAP plot (A) shows subsets of tanycytes, characterised according to previously-described makers (B). C. Violin plots show differential gene expression of selective genes in tanycyte subsets in NrCAM KO compared to wildtype hypothalamus.

Similar to the immunohistochemical and *in situ* analyses, we detected subtle changes in gene expression. As expected, we observed a decrease in *NrCAM* in KO compared to wildtype control mice in each tanycyte subset (Fig 6C) and a decrease in *Nestin* and *Rax* (Fig 6C), adding weight to the idea that tanycytes are reduced in the NrCAM KO mouse. We also observed reduced levels of *Hes1* and *Prdx6* - both previously implicated in regulating the balance of radial glial-like neural stem cells and neurons elsewhere in the CNS (Park et al.; Bai et al., 2007; Bosze et al., 2020) and reduced levels of *metr*, a gene implicated in astrocyte differentiation (Nishino et al., 2004) (Table 1). Finally, we observed lower levels of the tanycyte-enriched genes, *Ndn*, *Crym* and *Gpr50*, previously shown to regulate thyroid hormone signaling (Fig 6C; Table 1) (Bechtold et al., 2012; Hasegawa et al., 2012; Kinney and Bloch, 2021). At the same time, we detected a small increase in glial genes, such as *Slc1a4, Slc1a2* (solute carrier family 1 member), *Ntsr2* (neurotensin receptor 2) and *Mal* (Fig 6C, Table 1).

Previous lineage-tracing studies, showing that tanycytes can give rise to astrocytes (Lee et al., 2012; Haan et al., 2013; Robins et al., 2013; Yoo et al., 2021), and the reduced numbers of GFAP-expressing astrocyte cells in differentiated neurospheres obtained from NrCAM KO compared to wildtype mice (Fig 6C, I) prompted us to perform further analyses of astrocytes from wildtype and NrCAM KO littermates. We observed marked molecular changes in astrocytes between wildtype and NrCAM KO mice (Supp Fig S6). No NrCAM expression was detected and the classic astrocyte-enriched markers, *Ntsr2, Agt* (angiotensinogen), *Mt2* (metallothionein2), *Slc1a3* (solute carrier family 1 member 3), and *Gja1* (gap junction protein Connexin 43) were all expressed at lower levels in NrCAM KO compared to wildtype mice (Supp Fig S6, Table 2). *Htra1* - which encodes a serine protease that regulates TGFbeta and the availability of insulin-like growth factors - and has been not previously identified as an astrocyte-enriched marker - was also reduced. At the same time, other genes were upregulated, including *Vimentin* (*Vim* - a tanycyte-enriched intermediate filament-encoding gene), *Secretogranin* (*Scg2* involved in the packaging or sorting of peptide hormones and neuropeptides into secretory vesicles) and *Pfdn2* (Supp Fig S6, Table 2). In summary, scRNA seq analysis shows that astrocytic markers are reduced in the NrCAM KO mouse.

Finally, we interrogated hypothalamic neurons. Distinct subclasses could be identified, based on known molecular markers (Fig 7A). Comparative analyses revealed a subtle decrease expression levels of GABAergic neuron-associated genes (*Penk, Gabra1, Gria2*) and GABAergic markers (*Slc32a1, Slc17a6*) in NrCAM KO mice, relative to wildtype littermates (Fig 7B, Table 3). Additionally, we detected a small reduction in Thyroid releasing hormone (*Trh*) (Fig 7B, Table 3). However, the overall profile of neurons/neuronally-expressed genes was similar in mutant and wildtype mice. The decrease in tanycytes and astrocytes in NrCAM KO mice, therefore, does not lead to easily-detectable changes in hypothalamic neuronal profiles.

**Fig. 7.**
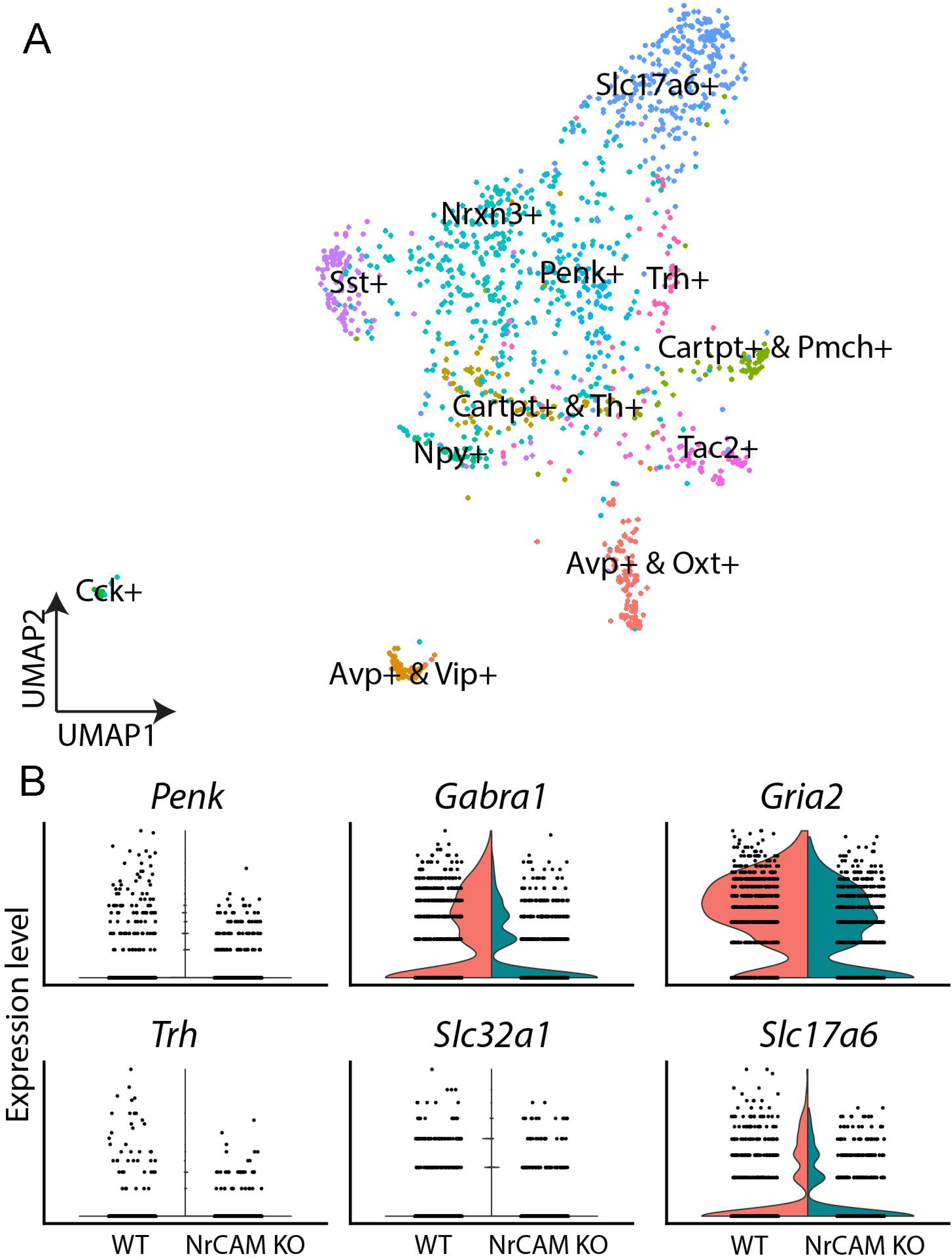
scRNA seq of neurons from wildtype and NrCAM KO mice. A. UMAP plot shows subsets of neurons, characterised according to previously-described makers. B. Violin plots show differential gene expression of selective genes in NrCAM KO compared to wildtype hypothalamus.

### Behavioural assays

The scRNA seq experiments suggest, nonetheless, that genes that govern thyroid hormone activity (*Trh, Ndn, Crym, Gpr50*) are subtly reduced in the NrCAM KO mice. Given the central role of thyroid hormone in metabolism, we therefore investigated whole body physiology in NrCAM KO mice versus wildtype (littermate) control animals. Weight at birth was similar in NrCAM KO and wildtype mice, and in post-weaned juvenile mice (weeks 5 and 6) fed standard chow, food intake was similar (Supp Fig S8). However, thereafter, in mice fed standard chow, bodyweight in NrCAM KO was significantly reduced, as was food intake (Fig 8A, B). Interestingly, in these animals, the respiratory exchange ratio (RER) was significantly reduced during the light phase (Fig 8C: two-way ANOVA time x treatment interaction p < 0.05). Oxygen consumption and total activity were unaffected in NrCAM KO animals (Fig 8D, E).

**Fig. 8.**
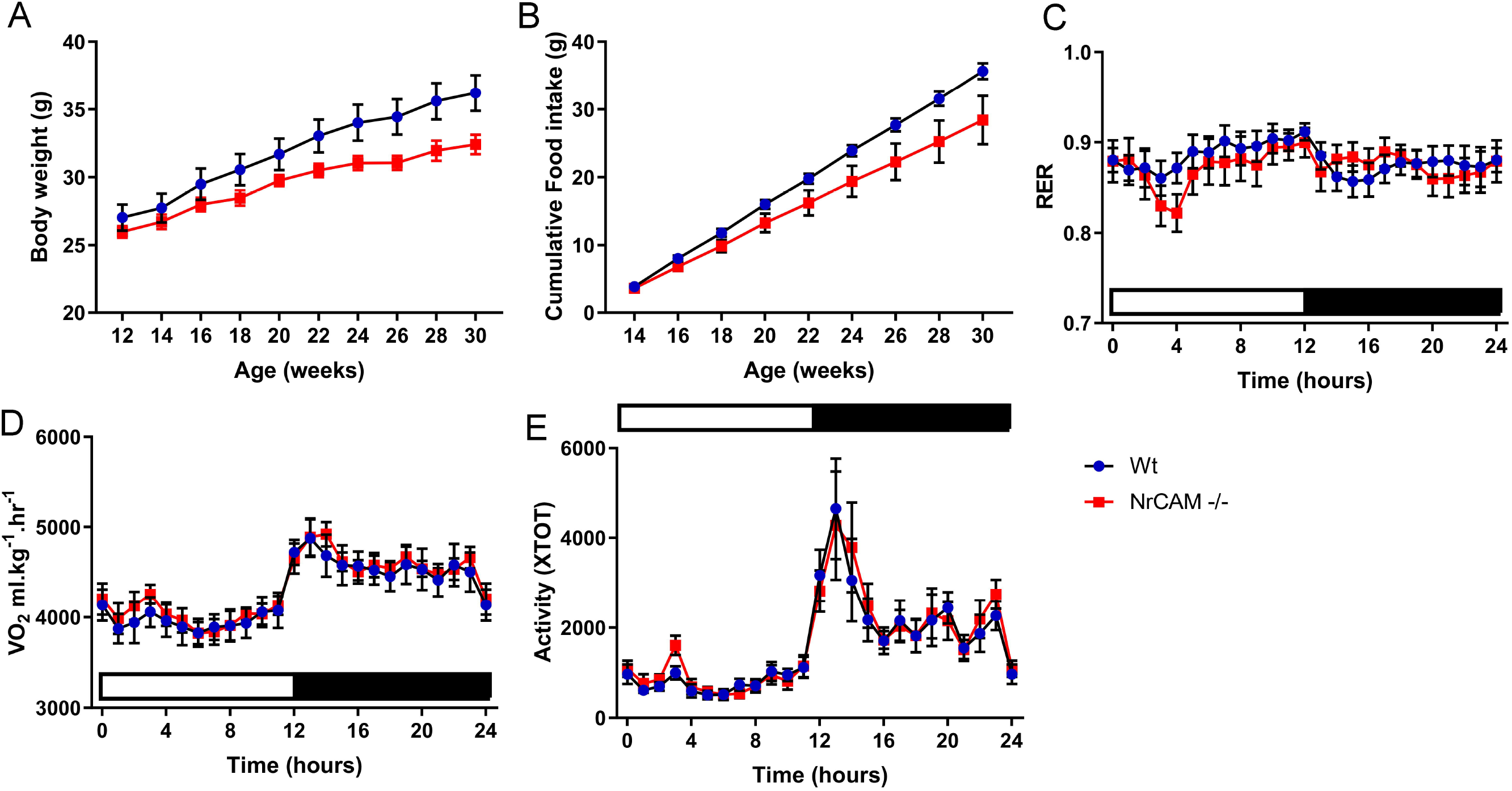
Body weight and food intake are reduced in adult NrCAM KO mice. NrCAM KO reduces body weight and food intake. (A) Body weight and (B) Cumulative food intake of wildtype and NrCAM KO mice in adulthood. (C) Respiratory exchange ratio (RER); (D) Oxygen consumption and (E) Activity of NrCAM KO mice versus wildtype littermate control animals (n = 8 per genotype). Values are mean ± SEM.

## Discussion

This study provides evidence that the neural cell adhesion molecule, NrCAM, regulates tanycytes that line the 3V of the hypothalamus. In wildtype mice, NrCAM is detected on hypothalamic embryonic radial glia and is then maintained at high levels on radial glia-derived tanycytes in the postnatal/adult animal. NrCAM KO mice, in which *NrCAM* RNA is reduced and NrCAM protein cannot be detected, have significantly fewer tanycytes than their wildtype siblings. Previous studies have suggested that tanycytes are stem and progenitor cells within a hypothalamic stem cell niche (Robins et al., 2013; Yoo et al., 2021), and that Rax, Lhx2 and Fgf10 play ongoing roles in late embryonic/postnatal life to support tanycyte differentiation and proliferation (Haan et al., 2013; Robins et al., 2013; Miranda-Angulo et al., 2014; Salvatierra et al., 2014). Our study adds to this, providing evidence that NrCAM supports cellular homeostasis within this niche.

The loss of NrCAM does not appear to interfere with early hypothalamic specification or growth: at E16, hypothalamic length and morphology, and expression domains of the transcription factors *Rax* and Six3, and the signalling ligands, *Fgf10* and *Shh* – all of which regulate tuberal hypothalamic progenitor domains (Placzek et al., 2020) – are indistinguishable in WT and NrCAM KO siblings. Likewise, the transcription factor, Lhx2 – also known to direct tanycyte differentiation (Salvatierra et al., 2014) – and Nestin - are indistinguishable at E16 in most of the hypothalamus in wildtype and NrCAK KO siblings. However, we detect a small but significant reduction in the density of Lhx2 and Nestinpositive radial glial cells in a limited spatial domain around the emerging ME. The reduction in radial glia in this region appears to prefigure a small but significant increase in embryonic TH-positive neurons in NrCAM mice compared to wildtype siblings – again, restricted to ME regions. The first manifestation of loss of NrCAM, therefore, appears to be a subtle imbalance, in late embryonic stages, in the numbers of radial glia and TH-positive Arc neurons.

By adulthood (8-10 weeks), there are significantly fewer tancytes throughout the hypothalamus in NrCAM mice compared to wildtype siblings. This is apparent through reduced numbers of Six3-positive VZ cells, a thinning of the VZ, reduced density of Nestin- and GFAP-positive tanycyte processes and a reduction in tanycyte-enriched genes, including *Hes1, Rax* and *Lhx2* (Shimogori et al., 2010; Lee et al., 2012; Salvatierra et al., 2014). ScRNA seq analysis of the tanycytic fraction demonstrates that the reduction in tanycyte-enriched genes is accompanied by an increase in glial markers. At the same time, scRNA seq demonstrates a marked reduction - in the astrocyte fraction - of astrocyte-enriched genes. Previous genetic lineage-tracing studies have demonstrated that adult tanycytes give rise to hypothalamic astrocytes (Lee et al., 2012; Haan et al., 2013; Robins et al., 2013; Yoo et al., 2021). Because NrCAM is expressed in astrocytes, we cannot exclude that the reduction in astrocyte-enriched genes is a direct consequence of the loss of NrCAM in astrocytes. However, an alternate interpretation is that the reduction in astrocytes is, as least in part, a consequence of a reduction in astrocyte-generating tanycytes. Support for this idea comes through neurospherogenic assays: under differentiation conditions, neurospheres from NrCAM KO hypothalamic tissue generate smaller numbers of GFAP-positive astrocytes than do neurospheres from wildtype siblings. Potentially, tanycytes and hypothalamic astrocytes are transcriptionally similar cell types, whose balance is disrupted by loss of NrCAM. An intriguing possibility is that tanycytes and astrocytes are similar cell states, as is the case in other brain niches, (Doetsch et al., 1999)9; (Cebrian-Silla et al., 2021). scRNA seq show changes, in NrCAM KO tanycytes, in the expression of *Metr, Ndn* and *Ntsr1* - all previously shown to govern neural stem cell proliferation and/or the differentiation of neural stem cells to astrocytes (Nishino et al., 2004; Huang et al., 2013; Ando et al., 2019) – raising the possibility that these may be part of a regulatory network that governs tanycyte and astrocyte states in the hypothalamus.

Our analyses extend previous characterisation of wildtype hypothalamic neurospheres (Robins et al., 2013) and reveals that, under non-differentiating conditions, they maintain expression of the regional stem/progenitor markers, Nkx2-1 and Six3, and also express NrCAM. Hypothalamic neurospheres derived from NrCAM KO mice maintain regional markers, but show decreased proliferation/ability to passage. The different behaviour of neurospheres from NrCAM and wildtype mice indicates a direct role for NrCAM in tanycyte regulation. Intriguingly, the NrCAM KO neurospheres follow a culture pattern that is similar to progenitor-like tanycytes, rather than stem-like tanycytes (Robins et al., 2013) – an initial increase in neurosphere number, followed by a decrease and failure to culture beyond late passage. Thus, NrCAM KO may directly affect the behaviour of stemlike tanycytes. Certainly in the intact NrCAM KO hypothalamus we detect a significant reduction in tanycytes in the region occupied by α2 (stem-like) tanycytes, as judged by the low numbers of GFAP-positive processes and Lhx2-positive nuclei. In other regions of the nervous system, NrCAM has been implicated in the control of progenitor cell proliferation. In the cerebellum, NrCAM was shown to be required to modulate granule neuron progenitor proliferation in response to SHH (Xenaki et al., 2011) and loss of NrCAM together withs its sister protein L1CAM leads to severe cerebellar hypoplasia (Sakurai et al 2001). More recently, NrCAM has been shown to be upregulated in androgen-treated proliferating neural stem cells (Quartier et al., 2018).

Numerous studies have shown that tanycytes can be stimulated to neurogenesis, but a key unknown question is whether tanycytes can give rise to hypothalamic neurons in the unchallenged animal. Our studies raise the possibility that tanycytes may generate TH-positive Arc neurons: we detect a small, but significant decrease in the adult, which contrasts with the increase in TH-positive neurons detected in the embryo. Further, we can detect the generation of TH-positive neurons, both in differentiating neurospheres and in an acute assay that monitors the behaviour of hypothalamic VZ cells. In each assay, significantly fewer TH-positive neurons are generated from NrCAM-KO tissue, compared to wildtype tissue. We did not, however, detect a decrease in *TH* in the scRNA seq analyses; potentially the low numbers of TH-positive Arc neurons are masked by the TH-positive neurons of the zona incerta, whose numbers are unaffected by loss of NrCAM. The scRNA-seq analysis does, however, show other subtle alterations in neuronal gene expression profiles, namely a reduction in GABAergic markers and Trh. Although we detect no NrCAM expression in neurons, the scRNA seq demonstrates neuronal expression of *NrCAM*, making it difficult to judge whether the alterations are a direct consequence of loss of NrCAM in neurons or an indirect effect due to a reduction in tanycytes.

The reduction in *Trh* expression in neurons, and reduction in tanycytic *Crym* - which encodes a thyroid hormone binding protein, *Gpr50* – is likewise involved in thyroid hormone metabolism, and *Ndn*, which encodes the Prader-Willi syndrome-deleted gene, *Necdin*, and modulates the thyroid axis (Hasegawa et al., 2012; Robins et al., 2013), raises the possibility that NrCAM KO animals may show alterations in metabolism. Indeed, from 8 weeks, NrCAM KO animals have lower body weight than wildtype littermate controls, and significantly reduced food intake. This subtle change appears to be a consequence of reduced food intake during the light phase, as the RER is significantly lower, suggesting increased fat oxidation. No change in energy expenditure or activity was apparent.

Finally, our systematic analysis demonstrates that as tanycytes develop from embryonic radial glia, they diversify along the A-P axis, and project, not only to the Arc and VMN but also to the periventricular nucleus (posterior part) (PVp). In contrast to tanycytes that project to the Arc and VMN, *Six3, Fgf10* and GFAP are either not detected or show weak expression in tanycytes that project to the PVp. Future studies are needed to determine the functional significance of this difference.

## Author contributions

A.F. and M.P. conceived the study. A.M., K.C., Sa.Br., I.S. performed in vivo characterisation. Sa.Br., K.C. A.M, S.R. M.P performed neurospherogenic and organotypic assays. Se.Bl and D.W.K. generated and analyzed scRNA-Seq data. I.S., J.E.L., G.D., C.M., M.T. and F.E. performed physiological assays. M.P.drafted the manuscript. Se.Bl., A.F., D.W.K., K.C., I.S., Sa.Br., J.E.L. and F.E. edited the manuscript.

## Acknowledgments

We thank Transcriptomics and Deep Sequencing Core (Johns Hopkins) for sequencing of scRNA-Seq libraries. This work was supported by the Wellcome Trust (212247/Z/18/Z) to MP, NIH (R01DK108230 and R01MH126676) to SB, the Maryland Stem Cell Research Fund (2019-MSCRFF-5124) to DWK and British Society for Neuroscience to JEL.

## Supplementary Figure Legends

**Fig S1: Selective expression of regional markers in neurospheres cultured from different regions of the CNS**

A, D, G. Coronal/transverse sections through 8-10 week adult tuberal hypothalamus (A), lateral ventricle of the brain (D) and spinal cord (G), after in situ hybridisation to detect *Six3*. Boxed regions shown in high power in (A’,D’,G’). Arrow in F’ points to weak expression of *Six3* in the subventricular zone (SVZ). Dotted outline in G’ shows ependymal region.

B-C, E-F, H-I. Representative examples of neurospheres cultured under non-differentiated conditions, analysed by in situ hybridisation for *Six3* expression (2 from each region to show range). Only hypothalamic neurospheres show strong *Six3* expression.

J-U. Representative examples of neurospheres from hypothalamus, SVZ and spinal cord ependymal region, cultured under non-differentiated conditions, analysed by immunohistochemistry. Neurospheres from all regions express the pan-neural stem cell markers, Sox2 and Vimentin, and proliferate (Ki67-positive), but only those from spinal regions express Nkx6-1.

**Fig S2: Comparative analyses in wildtype and NrCAM KO mice**

A-E Coronal sections through wildtype (A, C, E, F) or NrCAM KO (B, D, G, H) mice. (A-D) show sections at level of M, immunolabellled with anti-NrCAM or anti-GnRH Ab. (E-H) show serial coronal sections through AME, ME and posterior regions of the hypothalamus. (n=8 mice/genotype: analysed for NrCAM (n=5); GnRH, NPY and *Shh* (n=3). Arrowheads in (F’, G’) point to Shh midline cells. Scale bars: 100μm

**Fig S3: A13 TH-positive cells are unchanged in the NrCAM KO mouse**

Left hand panels: Immunolabelling of a coronal section taken through the AME, showing the dorsal zona incerta (A13) population in wildtype and NrCAM KO mouse. Scale bars: 100μm

Right hand panels: Quantification of TH-positive A13 cells (3 pairs of wildtype and NrCAM KO adult mice). Each icon represents a single measurement. Bars show 1 SD either side of the mean value. Analysis by unpaired t-test showed no significant difference of TH^+^ cells in dorsal AME Zona Incerta (A13) population between wild type and NrCAM KO adult mice (p=0.2543).

**Figure S4. Tanycyte progenitor markers at E16**

A-E. Representative serial coronal sections through AME, ME and posterior regions of the hypothalamus of a wildtype (A, B, D) or NrCAM KO mouse (C, E), analysed by immunohistochemistry to detect NrCAM and Nestin (A) or by *in situ* hybridisation to detect *Shh* and *Fgf10*. n=3 mice/condition. Scale bars: 100μm.

**Figure S5 scRNA seq analysis of the hypothalamus**

A. UMAP plots showing detected cluster

B. Violin plots showing Nrcam expression across cell types

**Figure S6 scRNA seq of astrocytes from wildtype and NrCAM KO mice**

Violin plots show differential gene expression of selective genes in astrocytes from NrCAM KO compared to wildtype hypothalamus.

**Fig. S7 Body weight and food intake in newborn-juvenile NrCAM KO and wildtype mice**

(A) Body weight in 3 pups of each genotype; (B) Average food intake of wildtype and NrCAM KO mice at 5 and 6 weeks (n = 6 per genotype). Values are mean ± SEM. Red - NrCAM KO; black - wildtype.

## Methods

### Animals

For molecular and neurospherogenic assays, adult mice were taken at 8-12 weeks, and embryonic mice at E16. NrCAM knock-out (KO) (Nrcam^tm1Gmt^/Nrcam^tm1Gmt^) transgenic mice, described elsewhere (Sakurai et al., 2001). Wild type and NrCAM KO siblings were bred within an NrCAM^+/-^ colony back-crossed to C57BL/6JOlaHsd for >10 generations. For molecular and neurospherogenic analyses, mice were kept in standard conditions for the duration of the study (18% protein rodent diet). Mice were housed in a centralized pathogen-free facility on a 12 hour light/dark cycle, at 19-23°C with 55% (±10%) humidity and 15-20 air changes per hour. For adult physiological studies, mice were re-derived at the University of Nottingham. Climates at both facilities were similar. All studies and procedures were conducted according to the UK Animals (Scientific Procedures) Act 1986/EU Directive 2010/63/EU, and were approved by the University of Sheffield Local Ethical Review committee (Licence 40/3742 to AF) and the University of Nottingham Local Ethical Review committee (Licence PFBB5B31F to FE).

### Tissue processing, immunohistochemistry and in situ hybridisation

Adult mice were anaesthetised by inhalation of isoflurane anaesthetic (B506; Abbott) before transcardial perfusion with 4% paraformaldehyde in 0.12 M phosphate buffer (4% PFA PB). Brains were dissected in ice cold Leibovitz’s 15 (L-15) media (11415-049; GIBCO), postfixed in 4% PFA PB at 4°C overnight, then transferred to 30% sucrose (S0389; Sigma), rocking at 4°C for 2-3 days until they sank. Prior to mounting, the anterior forebrain, hindbrain, and cerebellum were removed. The remaining medial portion of the brain containing the hypothalamus was orientated in OCT (361603E; VWR International) and frozen on dry ice before storage at −80°C.

Embryonic mice were collected in accordance with Schedule 1 methodology. Intact skulls were fixed in 4% PFA PB overnight at 4°C, brains removed and postfixed in 4% PFA PB for a further 3 hours at 4°C before transfer to 30% sucrose, rocking overnight. Brains were embedded in OCT, frozen on dry ice and transferred to −80°C for storage.

Sequential serial 15 μm-thick coronal sections using a cryostat (Leica Biosystems). For adults, sections were taken over a ~1500 μm length, covering Allen Brain Atlas sections 65-80. For embryos, sections were taken over a ~700 μm length, centred around the visibly obvious median eminence. For immunohistochemistry, serial floating sections were collected in PBS. For *in situ* hybridisation, serial sections were collected directly onto Superfrost Plus slides (Fisher Scientific).

For immunohistochemical analysis, adult samples were rinsed in PBS, then incubated in Citrate Buffer (10 mM Citric Acid (277804I; BDH), 0.05% Tween 20 (P9416; Sigma), pH 6.0) at 90°C for 1 hour for antigen retrieval. Sections were cooled for 20 minutes, rinsed in PBS and blocked in PBS with 0.5% TritonX-100 (T8787; Sigma) and 5% heat-inactivated goat serum (16210-072; GIBCO). Embryonic samples were rinsed in PBS and directly blocked in PBS with 0.1% TritonX-100 and 1% heat-inactivated goat serum. After a 1 hour block at room temperature, samples were incubated with primary antibody diluted in blocking buffer at 4°C overnight, rinsed in PBS, then incubated with secondary antibody in blocking buffer for 1 hour at room temperature. For adult tissue, DAPI (D9542-10MG; Sigma) was added at 100 ng/mL to the secondary antibody solution. Floating sections were mounted onto Superfrost Plus slides by careful manipulation with forceps and fine paintbrushes in a small quantity of 0.5X PBS. Excess liquid was removed from slides and sections coverslipped in Vectashield antifade mounting medium (#H-1200, Vector

Laboratories) before imaging on a Zeiss Axioimager Z1 with Apotome2 attachment and viewed in the Zeiss Zen 2 image acquisition software. Views show maximum intensity projections.

Antibodies used were as follows: mouse anti-GFAP (1:50, cat no. 556330, BD Pharmigen); mouse anti-MAP2 (1:1000, cat no M9942, Sigma-Aldrich); mouse anti-Nestin (1:200, cat no. Ab6142, Abcam); rabbit anti-NPY (1:1000, cat no. 22940, Immunostar); rabbit anti-NrCAM (1:1000, gift of Lustig et al 2001); rabbit anti-Nkx2.1 (custom antibody, Ohyama et al., 2005); rabbit anti-phosphoH3 (1:1000, cat no. 06-570, Millipore); anti-mouse RIP (1:10 cat no. AB531796, Developmental Studies Hybridoma Bank (DSHB)); anti-rabbit Six3 (1:10,000, Eurogentec custom antibody; sequence RLQ-HQA-IGP-SGM-RSL-AEP-GC); mouse anti-Slc1a3 (Glast, 1:1000, Ab 49643, Abcam); anti-rabbit Sox2 (1:200, cat no. Ab97959 Abcam); anti-rabbit TH (1:1000, cat no. Ab152, EMD Millipore); mouse anti-Pax6 (1:50, DSHB); rabbit anti-GHRH (1:600, AB1715, Chemicon); mouse anti-Vimentin (1:200, Sigma); mouse anti-Ki67 (1:500, Ab15580, Abcam); mouse anti-Nkx6.1 (1:50, F55A10, DSHB); goat anti-rabbit Alexa 488 (1:00, cat no. A11034, Molecular Probes); goat antirabbit Alexa 594 (1:500, cat no. A11012, Molecular Probes); goat anti-mouse IgG Alexa 488 (1:500, cat no. A11001, Molecular Probes); goat anti-mouse IgG Alexa 594)1:500, cat no. A11005, Molecular Probes).

Chromogenic *in situ* hybridisation was performed as previously described (Miranda-Angulo et al., 2014). Briefly, 15 μm coronal sections were fixed with 4% paraformaldehyde, permeabilised in PBS with 0.1% TritonX-100, then acetylated (1.3% triethanolamine (90279; Sigma), 2.1M HCl (H/1150/PB17; Fisher Scientific) and 0.25% acetic anhydride (added last) (A6404; Sigma)). Sections were hybridized with digoxigenin-labeled probes at 68°C overnight. Unbound probes were washed out and sections were blocked with sheep serum followed by incubation with anti-digoxigenin antibodies conjugated to alkaline phosphatase (1:5000) overnight at 4°C. Nitro-blue tetrazolium (NBT) and 5-bromo, 4-chloro, 3-indolylphosphate (BCIP) were used as chromogenic substrates of alkaline phosphatase.

Colour development was continued for the same amount of time for wildtype and NrCAM KO sections.

### Neurospherogenic assays and organotypic cultures

Animals were sacrificed using a lethal dose of isoflurane anesthetic and cervical dislocation. Brains were removed into ice-cold L-15, and the hypothalamus was manually isolated.

Neurospherogenic assays were performed as previously described (Robins et al., 2013). Briefly, hypothalamic regions close to the VZ were isolated from adjacent parenchyma using tungsten needles, minced and enzymatically dissociated using TrypLE enzyme solution (12604013, Life Technologies) for 20 min at 37°C, then mechanically dissociated by trituration to single-cell suspension using 25-gauge and 30-gauge needles. Cells were plated in Corning Costar Ultra-Low attachment 24 well plates (3473 SLS) in DMEM:F12 (21331; GIBCO) supplemented with 100 U/mL penicillin/streptomycin (11578876: Fisher Scientific), 2 mM L-Glutamine (25030024; GIBCO), 1X B27 (17504-044; Life Technologies), 5 μg/ml heparin, 10 μM transferrin (T0665; Sigma), 20 nM progesterone (P7556; Sigma), 30 nM selenite (S5261; Sigma), 100 μM putrescine (P5780; Sigma), 50 ng/mL IGF-1 (I8779-50UG; Sigma), 20 ng/mL bFGF (13256029; Life Technologies), and 20 ng/mL EGF (PHG0314; Life Technologies). Primary neurosphere cultures were maintained at 37°C and 5% CO2 and assessed 7 days after plating. Neurospheres were passaged by incubating for 20min in TrypLE solution, before dissociating and plating as per the original conditions. From first passage onwards, neurospheres were fed every other day. Neurospheres were differentiated in Nunc Lab-Tek 8-well Permanox chamber slides (C7182; Sigma) coated with poly-D-lysine (P1024; Sigma) for 1 hour and fibronectin (10042682; Fisher Scientific) for 4 hours. Differentiation media was made to the same specification as neurosphere culture media but without IGF-1, EGF, reduced bFGF (10 ng/mL), and with B27 supplement replaced with B27 minus insulin (A1895601; Thermo Fisher). One neurosphere was added to each well and cultured for 7 or 14 days as specified. Differentiation cultures were fed every two days, before fixation and processing.

For organotypic assays, the hypothalamus was mounted coronally on a McIlwain tissue chopper and 100 μm-thick slices were obtained. Electrolytically-sharpened tungsten needles were used to isolate the VZ from tuberal slices. Explants were embedded in collagen, as described elsewhere (Placzek & Dale (1999) and cultured in 0.5 mL Opti-MEM (31985070; Fisher Scientific) with 100 U/mL penicillin/streptomycin (11578876; Fisher Scientific), 2 mM L-Glutamine (25030024; GIBCO), and 4% FBS (10042682; Fisher Scientific) (Placzek et al., 1993) supplemented with 3 nM Shh (1845-SH; R&D Systems), as used in Ohyama *et al* (2005) to promote hypothalamic neuron differentiation from prospective hypothalamic tissue. Following culture, explants were fixed in 4% PFA PB for 2 hours at 4°C, transferred to 30% sucrose overnight, then embedded in OCT for cryostat sectioning at 15 μm.

### Tanycyte process and cell counts

Nestin-positive process counts and TH-positive cell counts were performed blindly, on every alternate section (~30 sections in adult covering sections every 30 μm between Bregma −0.9 to −2.2/ every 30 μm over ABRA sections 67-77); 15 sections in embryo, centred around ME) in the regions covered by α2- and β1-tanycytes; GFAP-positive process counts and DAPI-positive measurements were performed on every alternate section over the length of the ME (ABRA sections 69-73). Every 5th section was analysed independently by a second individual. Processes were marked by eye with the ImageJ multipoint tool, where they first extend into the SVZ from the VZ. DAPI counts Process numbers were normalized to length, to obtain a measure of process density (processes per μm). All values are expressed as means ± *SD*.

For neurospherogenic assays, cells were counted in 20x random fields; in organotypic assays, cells were counted from each 15μm section taken through the entire explant.

### Quantification of chromogenic ISH signal

Quantification of *Rax*, *Fgf10*, and *Shh*-expression in the VZ was performed blindly, on every 5^th^ section (adult regions sampled over ABRA sections 67-77; embryonic sections sampled around ME). Length measurements were collected using the ImageJ segmented line tool. Signal intensity was obtained from a one-pixel line profile through the chromogenically labelled domain, and the mean gray value was measured. Intensity was normalised to a line profile through the adjacent non-labelled parenchyma. Isolated Shh or Rax-positive cells were quantified using the ImageJ multi-point tool.

### scRNA seq analysis

Hypothalamic tissue was manually dissected and transferred to Hibernate E medium (Cat No. HE500, BrainBits LLC). Tissue was enzymatically digested using Papain (2 mg/ml, Cat No. LS003119, Worthington Chemicals), and cells dissociated into single cells using fire-polished glass pipettes. Papain was neutralized with Hibernate-A medium with B27 and GlutaMAX (Kim et al., 2020). Cells were pelleted at 500g at 4°C, washed twice in 1 ml cold 1 x PBS, and then resuspended in 200 ul chilled 1 x PBS by gentle pipetting. Cells were then fixed with 800 μl cold 100% methanol as previously described (Alles et al., 2017). Cells were stored at −80°C until scRNA-Seq, then re-hydrated as previously described (Alles et al., 2017). Briefly, cells were washed twice (3000 x g for 5 min at 4°C) and resuspended in RNAse-free PBS with 1% BSA and 0.5 U/ul RNase inhibitor (Cat. N2615, Promega). Cells were then used for the 10x Genomics Chromium Single Cell System (10x Genomics, CA, USA) using V3.0 chemistry per manufacturer’s instruction, loading between 8,000 and 12,000 cells per run. Libraries were sequenced on Illumina NextSeq 500 with ~ 400 million reads per library. Sequenced files were processed through the CellRanger pipeline (v.3.10, 10x Genomics), and scRNA-Seq analysis was conducted as described previously (Kim et al., 2020)

### Physiological measurements

Bodyweight and food intake in the home cage were measured at two-week intervals. At approximately 2, 6 and 9 months of age energy expenditure parameters and feeding behaviour were measured using a comprehensive lab animal monitoring system (CLAMS: Linton Instrumentation, Linton, UK, and Columbus Instruments, Columbus, OH). This is comprised of 8 chambers in which mice were individually housed, with water bottles and food hoppers located at the centre of each chamber. Metabolic activity was measured in a variety of ways including oxygen consumption (VO_2_) and respiratory exchange ratio (RER; CO_2_ production/VO_2_). Locomotor activity was measured via infrared beams that cross each cage, monitoring movement along the x, y, and z axes. Locomotor activity was calculated as the number of infrared beam breaks recorded in 9-minute time intervals. Feeding behaviour of the animals was also monitored to establish total food intake and parameters such as meal size and duration and timing of each feeding bout. The diet used for this study consisted of standard laboratory chow comprising 19% extruded protein and 9% fat (Teklad 2019, Harlan, UK). All raw metabolic data were recorded in 9-minute intervals and collected from the CLAMS using OxyMax software (v4.2, Columbus Instruments). Male mice (8 NrCAM KO, 8 wild-type littermates) were placed in the CLAMS at approximately 2 months of age (mean= 8.6 weeks, range: 7.9-10.9 weeks), 6 months of age (mean=26.3 weeks, range: 24.9-28.9 weeks), and 9 months of age (mean= 36.3 weeks, range: 31.0-39.4 weeks). The mice were maintained on a 12 hour light/dark cycle, and were initially placed in the metabolic chambers during the light phase, and maintained for 48 hours. Data for the second 24 hour period were analysed to allow the mice time to acclimatise to the chambers.

### Statistical analyses

Analyses of studies with two groups (genotypes) were performed through two-tailed Student’s t-test (Microsoft Excel). Chromogenic signal measurements were statistically analysed in GraphPad Prism 7.04 using the Wilcoxon matched-pairs signed-rank test. Descriptive statistics (means ± SEM) were generated using GraphPad Prism (version 9.0, San Diego, CA, USA). Data were analysed using two-way ANOVA. No animals were excluded from the analyses. Statistical significance was considered at p < 0.05.

