## Supplementary figures and images for "The neural cell adhesion molecule NrCAM regulates development of hypothalamic tanycytes"

### Supp Fig 1

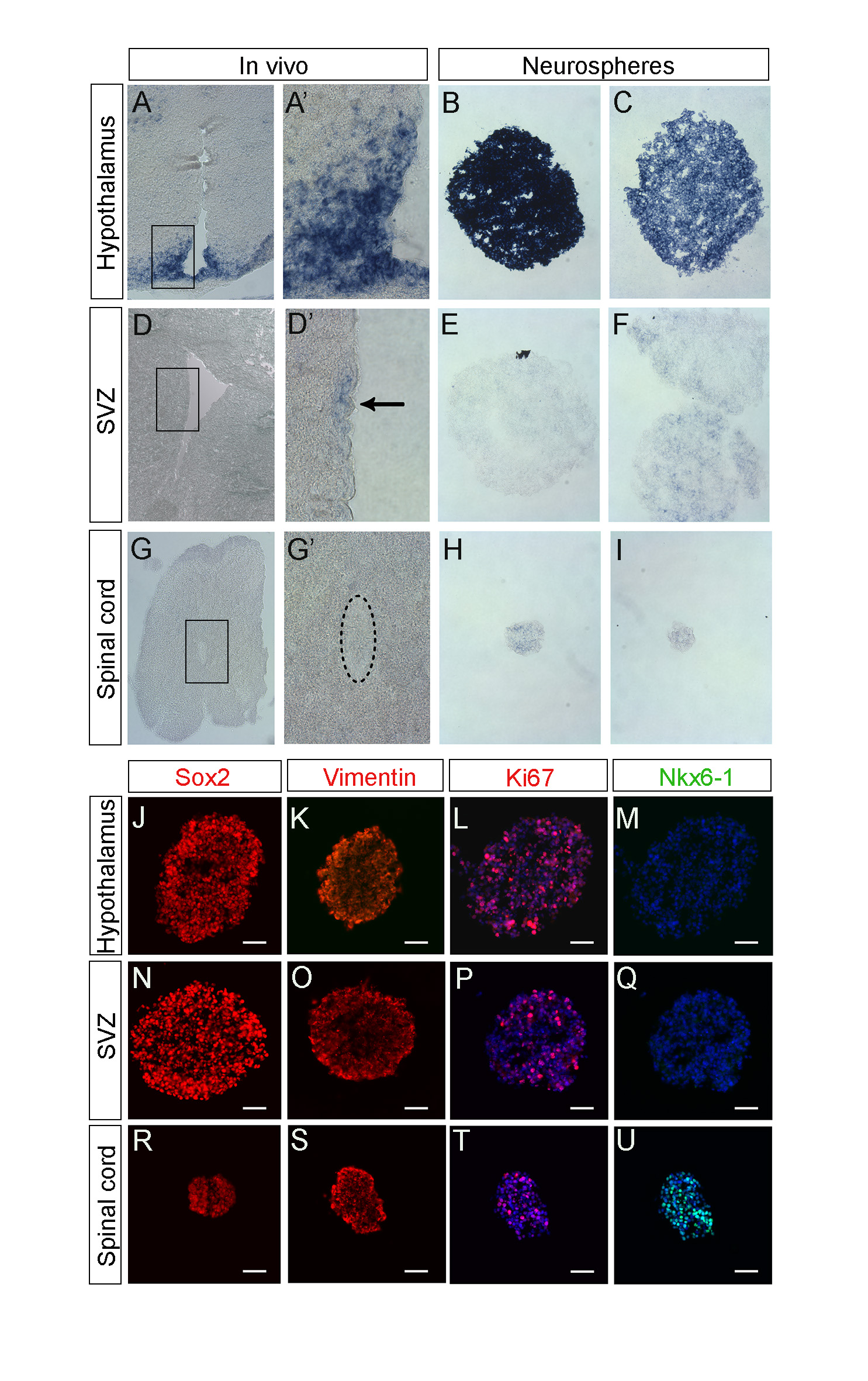

### Supp Fig 2

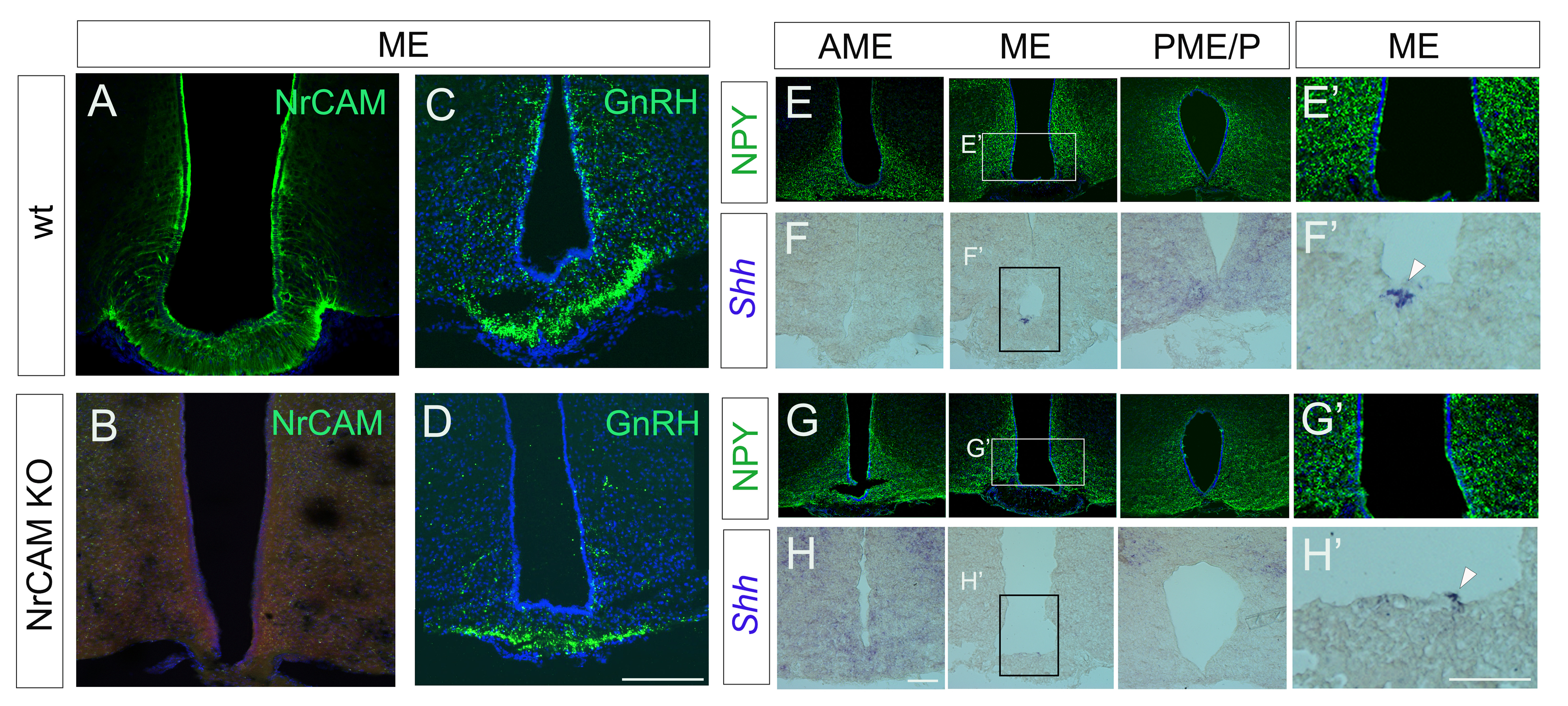

### Supp Fig 3

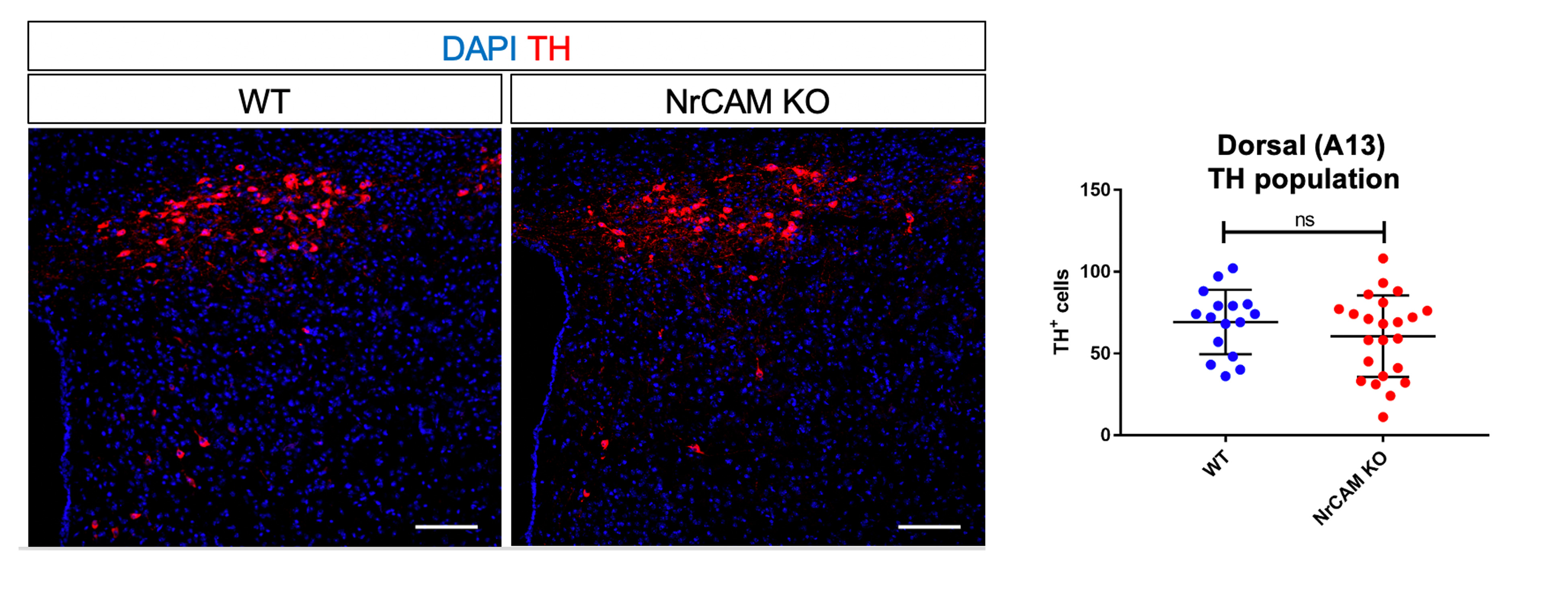

### Supp Fig 4

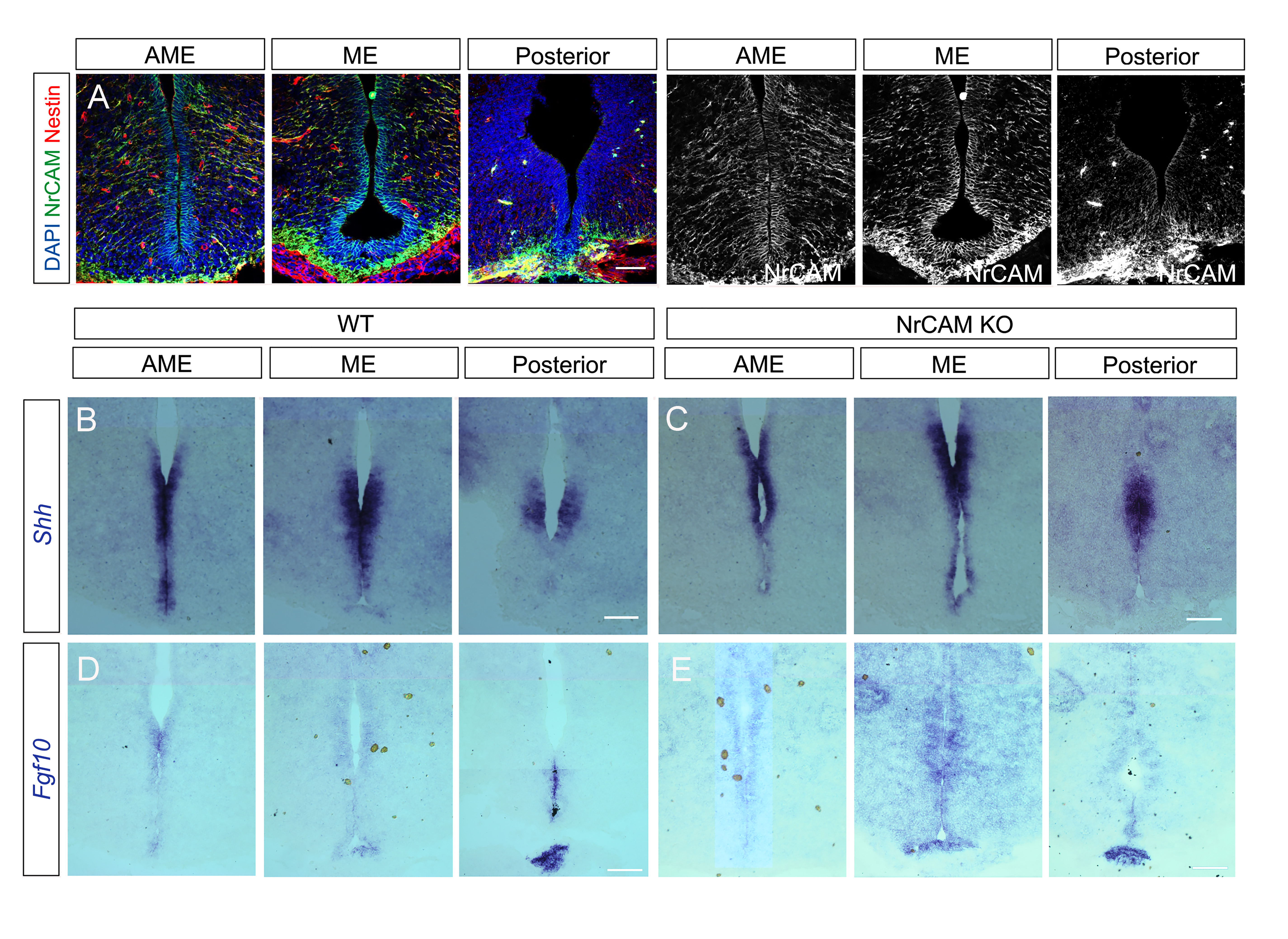

### Supp Fig 5

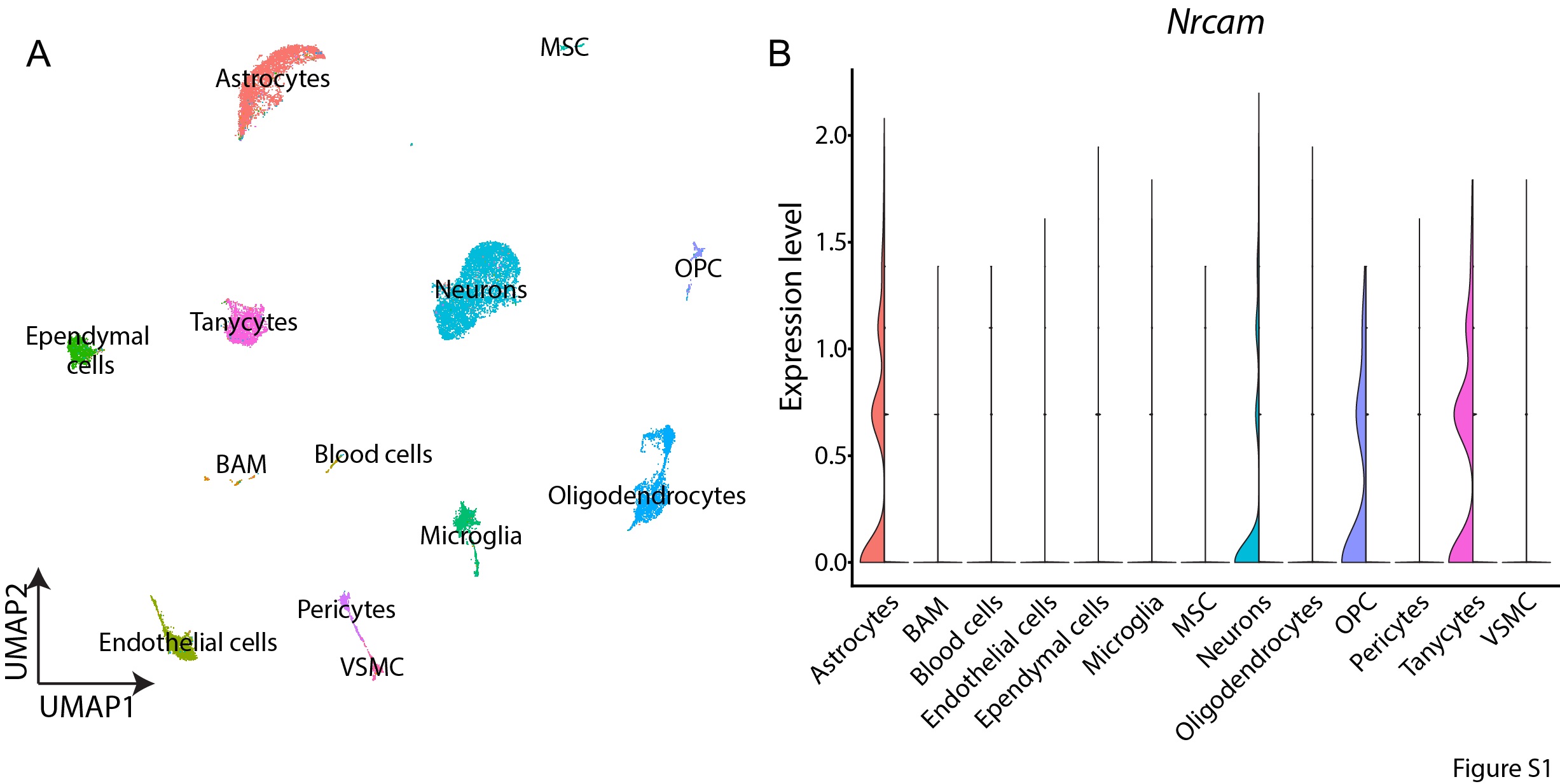

### Supp Fig 6

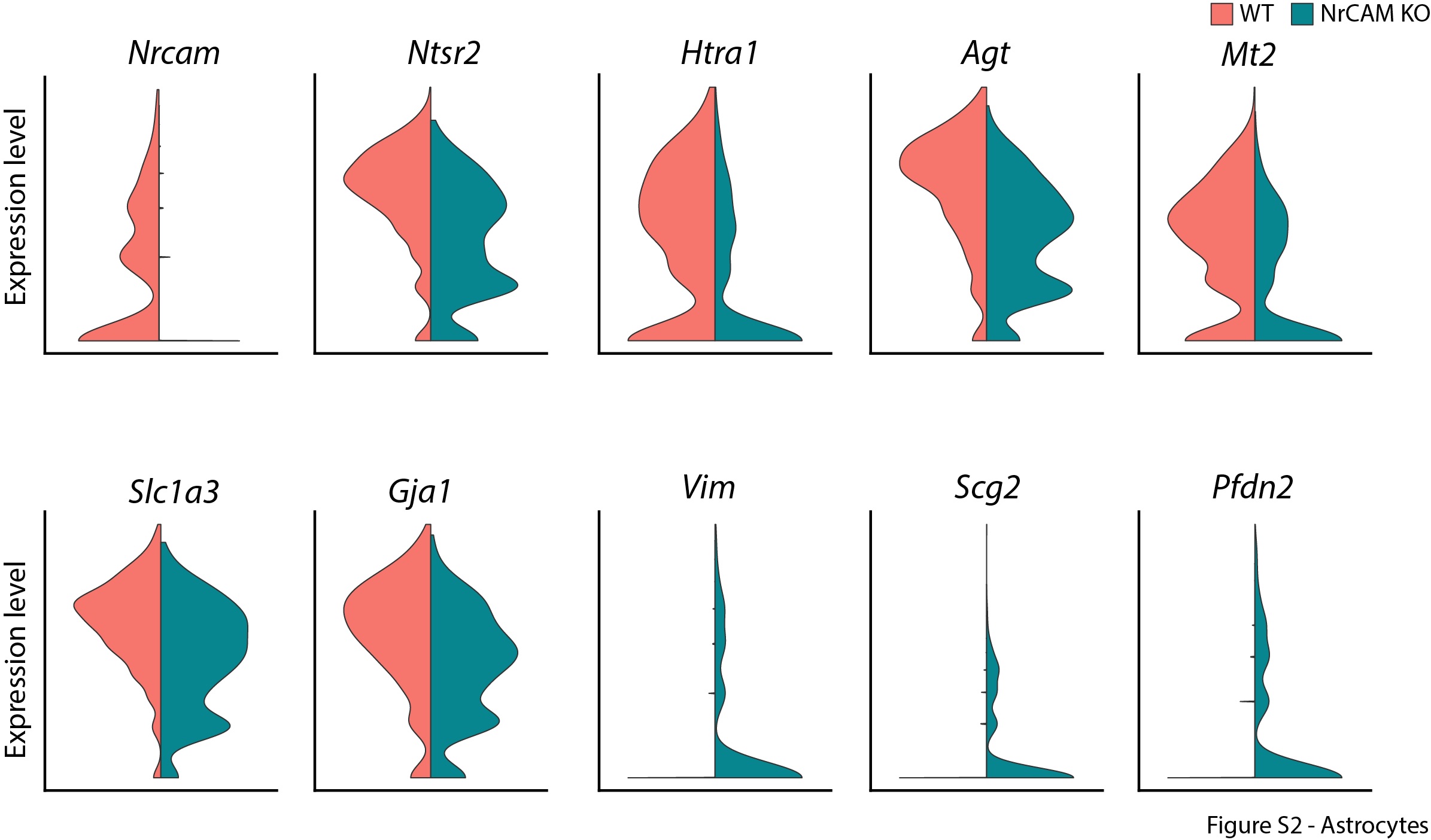

### Supp Fig 7

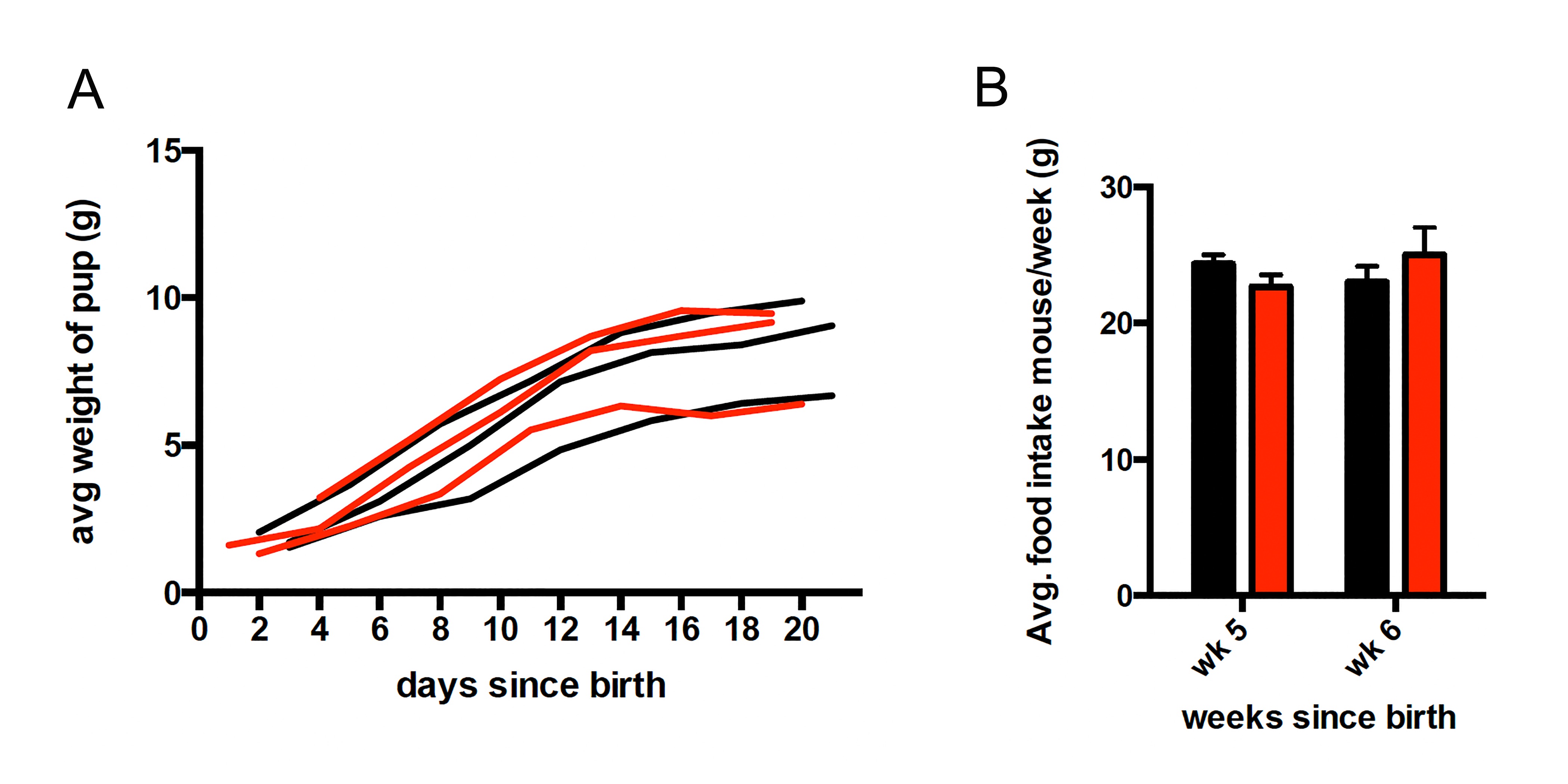
